# Rapid reduction in global chromatin loop size after acute STAG2 reconstitution in human cancer cells

**DOI:** 10.64898/2026.05.26.727913

**Authors:** Tianyi Yang, Jung-Sik Kim, Clara Mellows, Wanying Xu, Alvin Ya, Lisa Sadzewicz, Luke Tallon, Jann Sarkaria, Fulai Jin, Todd Waldman

## Abstract

Truncating mutations in the tumor suppressor *STAG2*, which encodes a component of the cohesin complex, are prevalent across diverse human cancers. Here, we report that acute reconstitution of physiological STAG2 levels in *STAG2*-mutant human glioblastoma multiforme cancer cells triggers a rapid reduction in the size of chromatin loops genome-wide. Despite this global change in chromatin loop size, early transcriptional responses to STAG2 restoration are limited to a small number of genes, most of which are induced by STAG2 reconstitution. Notably, the growth-suppressor EFEMP1 (Fibulin-3), a secreted glycoprotein that functions as an extracellular matrix-associated inhibitor of glioblastoma growth and invasion, was the only gene consistently induced across all experimental models. The most robust and conserved STAG2-induced genes all reside within intense chromatin loops whose anchors and overall intensity did not appear to change in response to STAG2 reconstitution. These findings suggest that inactivating mutations of STAG2 promote neoplastic transformation by alleviating a restriction on chromatin loop size, allowing for an expanded range of chromatin interactions that disrupts the maintenance of a tumor-suppressive transcriptome.

## INTRODUCTION

Inactivating somatic mutations in the tumor suppressor gene STAG2 are prevalent across a wide spectrum of human malignancies, including bladder cancer, glioblastoma multiforme (GBM), myeloid leukemias, and Ewing sarcoma (1,2,3). Because STAG2 is located on the X chromosome, it is functionally haploid in both males and females, the latter due to X-inactivation. Approximately 85% of tumor-derived STAG2 mutations are truncating, typically resulting in the unambiguous, complete loss of protein expression. Consistent with its role as a tumor suppressor, mice with targeted STAG2 inactivation are predisposed to myeloid neoplasia and exhibit increased aggressiveness in several solid tumor models (4,5,6,7,8).

The STAG2 protein is a component of cohesin, a ring-shaped multi-protein complex that mediates sister chromatid cohesion and generates the chromatin loops that underlie 3D genome organization (9,10). However, despite over a decade of research, the precise mechanisms by which STAG2 loss drives tumorigenesis remain elusive. Current evidence suggests that STAG2, and the cohesin complex more broadly, maintains stemness and regulates differentiation, particularly in myeloid lineages (4,5,11,12,13,14). However, generalizing these findings to solid tumors has proven difficult.

Mechanistic models of STAG2 tumor suppression focus on its role in 3D genome architecture. Consequently, much research has aimed to identify genes whose expression is directly controlled by STAG2-containing cohesin complexes. However, a major technical hurdle has been the inability to rapidly reconstitute physiological levels of wild-type STAG2 in mutant human cancer cells. This limitation has made it difficult to distinguish between the primary, direct effects of STAG2 and the indirect, secondary changes that accumulate in steady-state models.

To address this, we developed a STAG2-expressing adenovirus capable of rapidly restoring STAG2 to physiological levels in mutant cells. By interrogating chromatin architecture and gene expression immediately following STAG2 reconstitution, we found that STAG2 imposes a rapid and global constraint on chromatin loop size. We further identified a small, discrete set of rapid STAG2-responsive genes - most notably the growth suppressor EFEMP1 - that were robustly induced across all models. These findings suggest that STAG2 inactivation promotes neoplastic transformation by alleviating a restriction on the scale of loop extrusion, allowing for an expanded range of chromatin interactions that disrupts the maintenance of a tumor-suppressive transcriptome.

## RESULTS

### Acute physiological reconstitution of STAG2 in human GBM cell lines

To investigate the immediate consequences of STAG2 restoration, we developed a recombinant adenovirus expressing wild-type STAG2 (adeno-STAG2; ref. 15). We applied this adenovirus to two independent human GBM cell lines harboring endogenous *STAG2* mutations: H4 (*STAG2* 357N>fs; ref. 16) and MOG-G-UVW (*STAG2* 216R>STOP; ref. 17). We previously used gene editing to generate and study a permanently STAG2-corrected H4 derivative (1). The H4 model therefore provided a suitable experimental system for comparing the acute effects of STAG2 reconstitution (via adenoviral delivery) to the chronic effects observed after permanent gene correction.

In both models, STAG2 protein was restored to levels comparable to *STAG2* wild-type cells within 36 hours (Fig. 1A,B), with transduction rates exceeding 98% in both lines as confirmed by GFP flow cytometry. Interestingly, acute STAG2 reconstitution did not significantly alter cell proliferation or clonogenic survival compared to vector-only controls (Fig. S1). The absence of a clear phenotype allowed us to characterize the primary structural and transcriptional consequences of STAG2 restoration without the confounding effects of altered cell cycle kinetics or reduced viability.

**Figure 1.**
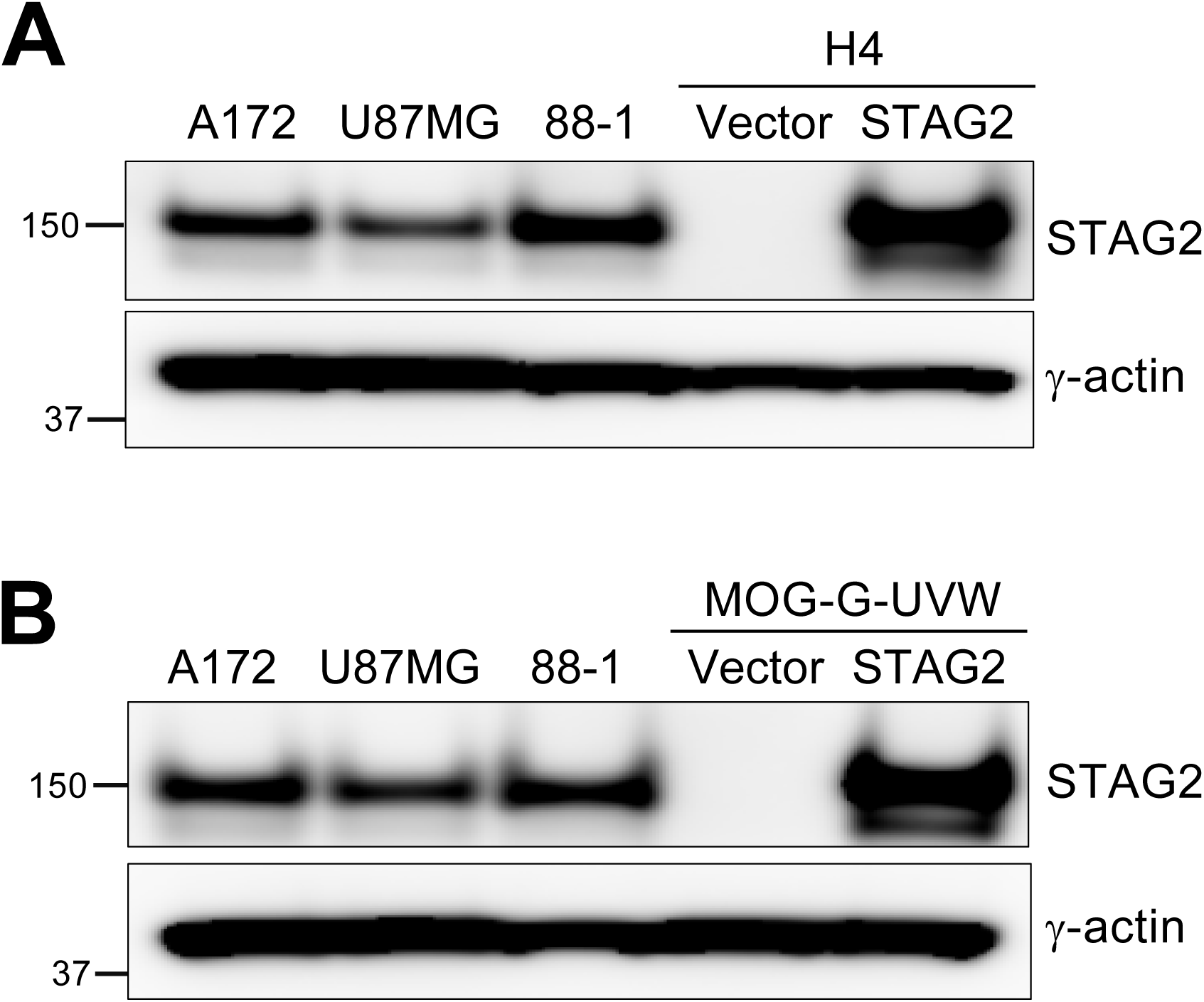
Reconstitution of STAG2 expression in human GBM cells. Western blot of STAG2 levels in (A) H4 and (B) MOG-G-UVW cells 36 h post-infection with control adenovirus (Vector) or Adeno-STAG2. STAG2 wild-type GBM lines A172, U87MG, and H4 88-1 STAG2 corrected cells are included to indicate levels of endogenous STAG2 expression in wild-type cells.

### STAG2 reconstitution triggers a rapid, global, and permanent reduction in chromatin loop size

Studies have indicated that STAG1-cohesin complexes possess greater chromatin residence time than STAG2-cohesin complexes, resulting in greater processivity and the extrusion of longer chromatin loops (18). We hypothesized that acutely restoring the STAG2/STAG1 balance in *STAG2*-mutant human cancer cells would impose a constraint on loop extrusion and result in a rapid reduction in chromatin loop size. To test this, we performed high-resolution Hi-C on H4 and MOG-G-UVW cells 36 hours post-infection with adeno-STAG2 or vector control (Table S1).

Following data processing with *HiCorr* to remove systematic biases and *DeepLoop* to enhance signal-to-noise ratio and resolve fine-scale interactions (19,20), we utilized Aggregate Peak Analysis (APA; ref. 21) to determine the effect of acute STAG2 reconstitution on chromatin loop size genome-wide. To quantify shifts in the loop landscape, we partitioned the 300,000 strongest chromatin loops into four size bins: 50-100 kb, 100-200 kb, 200-500 kb, and 500-2000 kb. As shown in Fig. 2A-B, STAG2 reconstitution resulted in an increase in the intensity of smaller loops (<200 kb) and a corresponding weakening of larger loops (>200 kb). This rapid reduction in loop size was conserved across both H4 and MOG-G-UVW models.

**Figure 2.**
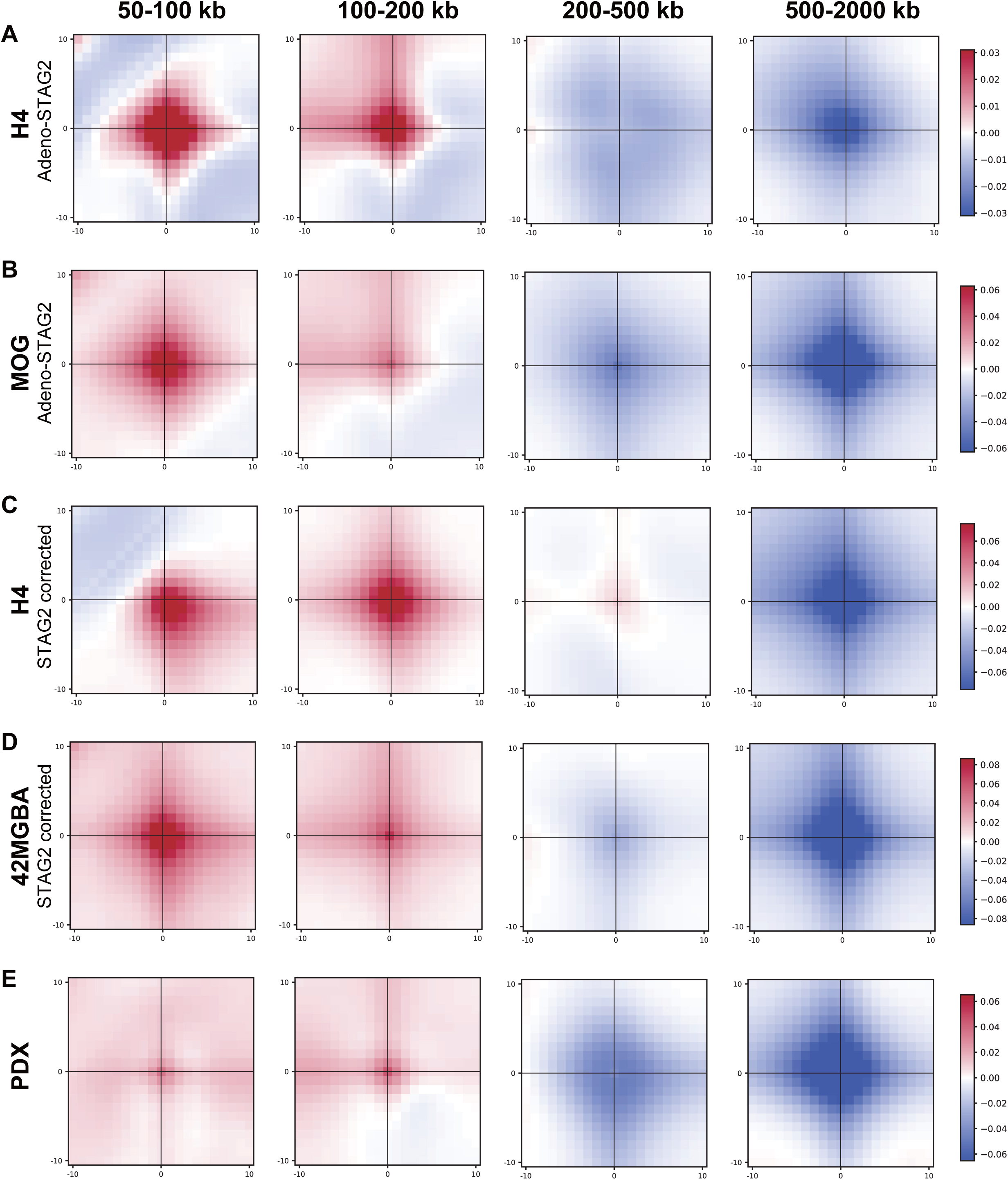
STAG2 reconstitution reduces chromatin loop size genome-wide. Subtraction aggregate peak analysis (APA) plots (STAG2-WT minus STAG2-mutant) showing normalized loop intensity for the top 300,000 loops. Data are categorized by genomic distance: 50-100 kb, 100-200 kb, 200-500 kb, and 500 kb-2 Mb. The central 10×10 pixel area (50 kb) represents the loop anchors. Red indicates increased loop intensity in STAG2-reconstituted cells; blue indicates increased intensity in STAG2-mutant cells. In these APA plots, center signal indicates equivalent loop size; enrichment in the upper-left quadrant signifies loop shortening in the WT relative to mutant, while the lower-right quadrant signifies loop lengthening. Panels show comparisons for: *A*, H4 cells infected with Adeno-STAG2 vs. Vector; *B*, MOG-G-UVW cells infected with Adeno-STAG2 vs. Vector; *C*, STAG2-corrected H4 clones (88-1) vs. parental STAG2-mutant H4 cells; *D*, STAG2-corrected 42MGBA clones (53-1) vs. parental STAG2-mutant 42MGBA cells; and *E*, STAG2-WT primary GBM xenografts (n=2) vs. STAG2-mutant xenografts (n=2).

To ensure this change was not a transient response to acute STAG2 expression, we performed an identical APA analysis to evaluate the loop landscape in two additional isogenic sets of STAG2-mutant human GBM cells and their permanently gene-corrected derivatives: H4 (clone 88-1) and 42MGBA (clone 53-1; ref. 22). Remarkably, the reduction in global chromatin loop size in these permanent models was virtually identical to that observed following acute STAG2 reconstitution (Fig. 2C,D), confirming that loop contraction is a stable and conserved feature of the STAG2-reconstituted state. To determine if this shift occurs in clinical specimens, we extended our APA analysis to high-resolution Hi-C data from four human GBM primary xenograft tumors (22) - two *STAG2* wild-type and two *STAG2*-mutant. Consistent with our cell line models, *STAG2*-mutant tumors exhibited larger chromatin loops than wild-type tumors (Fig. 2E). The persistence of this phenomenon across clinical tumor samples, three distinct GBM cell lines, and multiple methods of STAG2 restoration indicates that STAG2 serves as a fundamental and robust constraint on chromatin loop size.

### STAG2 reconstitution selectively strengthens smaller chromatin loops

To determine how STAG2 restoration impacts the strength of individual loops, we compared chromatin loop intensity in adeno-STAG2-infected cells and vector controls using regression analysis. This revealed that fewer than 1% of loops showed significant changes in intensity (Fig. 3A-D), and no loops exhibited an absolute dependence on the presence or absence of STAG2. Importantly, the number of altered intensity loops was substantially lower than previously observed in permanent gene-corrected models (22), suggesting that most changes in chromatin loop intensity in stable models reflect secondary, downstream effects. However, most loops that did change were conserved across both H4 and MOG-G-UVW lines and showed significant overlap with loops identified in our permanent H4 correction models (Fig. S2,S3).

**Figure 3.**
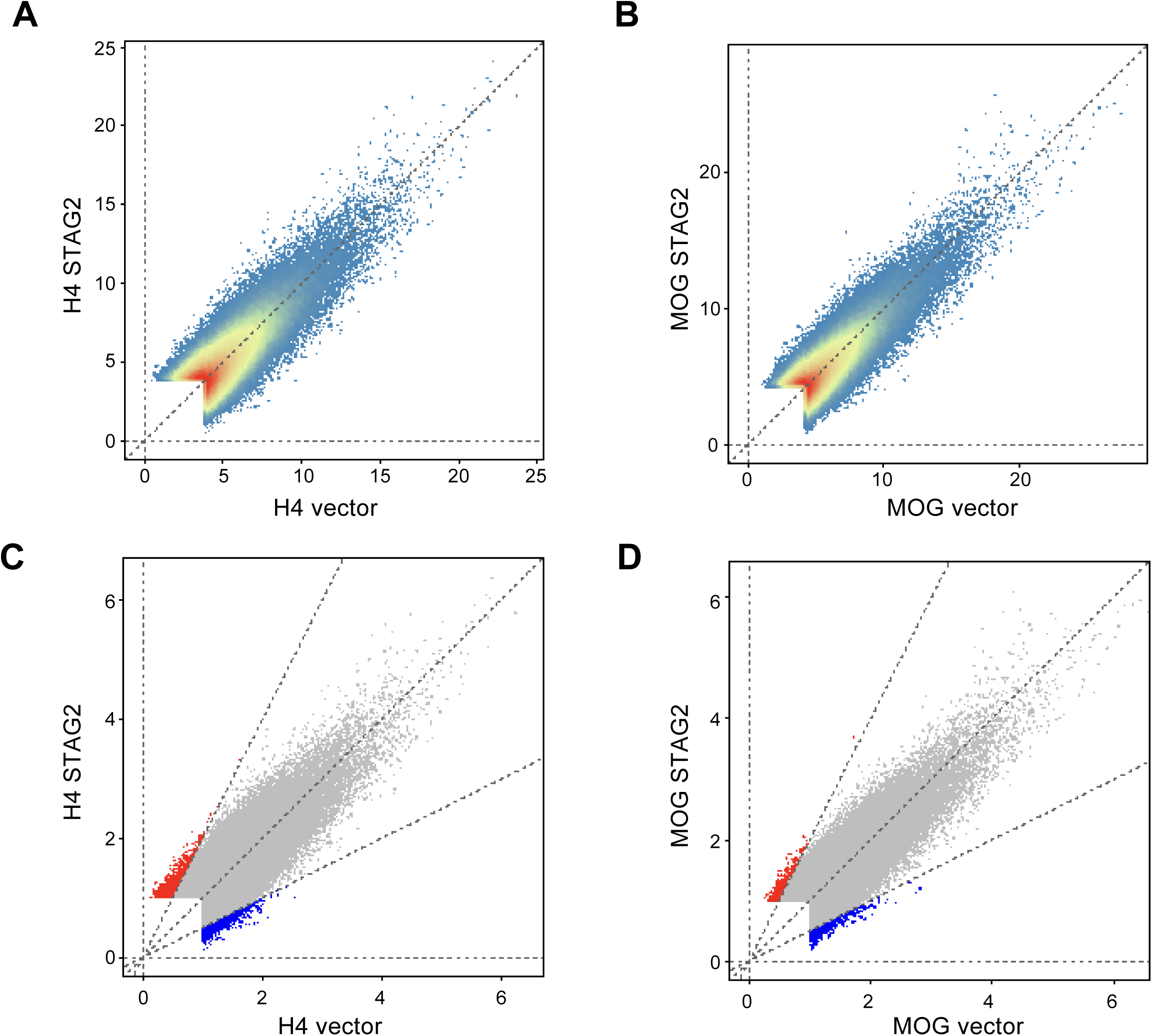
Impact of rapid STAG2 reconstitution on chromatin loop strength. *A*-*B*, Scatterplots showing the correlation of chromatin loop intensities between H4 (A) and MOG-G-UVW (B) cells 36 h post-infection with control vector or adeno-STAG2. Analysis of the strongest chromatin loops (∼3.8×10⁵ for H4; ∼3.7x 10⁵ for MOG-G-UVW) demonstrates a high level of correlation (R^2^ = 0.59 and 0.66, respectively). *C*-*D*, Re-plotting of data from (A) and (B) highlighting differentially regulated loops. Blue pixels indicate loops weakened by STAG2 reconstitution, and red pixels represent strengthened loops (FC>2).

An analysis of these altered intensity loops revealed a strong correlation with loop size. Loops strengthened by STAG2 were significantly smaller (median ∼200 kb) than the global average, while weakened loops tended to be larger (median ∼700 kb; Fig. 4). These findings indicate that STAG2 reconstitution preferentially strengthens small loops and weakens large loops, consistent with the lower processivity of STAG2-cohesin complexes.

**Figure 4.**
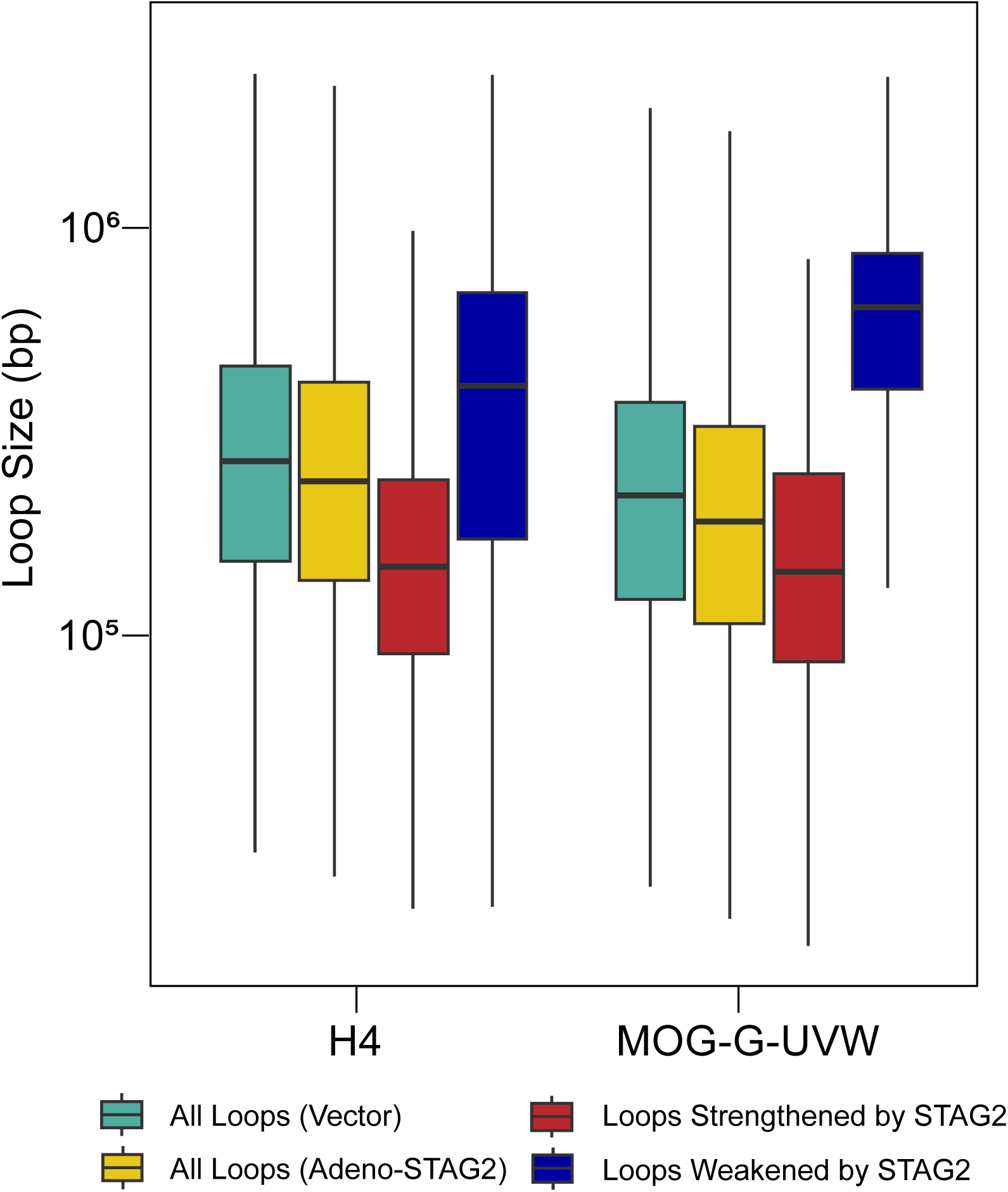
STAG2-intensified loops tend to be small. Histograms illustrate the distribution of chromatin loop sizes in H4 and MOG-G-UVW cells. Size distribution of the ∼300,000 strongest chromatin loops in cells infected with control vector (green) or adeno-STAG2 (yellow) confirms the reduction in overall loop size following STAG2 reconstitution. Further analysis of the size of loops whose intensity is regulated by STAG2 (depicted in Fig. 3C,D) reveals that loops strengthened by STAG2 (red) tend to be smaller, while loops weakened by STAG2 (blue) are larger.

### STAG2-mediated chromatin changes are restricted to loops

To determine if the reduction in loop size reflected a broader reorganization of the genome, we next assessed large-scale features including A/B compartments and Topologically Associating Domains (TADs). Acute STAG2 reconstitution had virtually no impact on A/B compartment profiles in either cell line (Fig. S4). This lack of immediate change contrasts with our previous findings in permanently corrected H4 cells (22), where 3% of compartments switched, suggesting that compartment switching is an indirect downstream consequence of STAG2 restoration. Similarly, rapid STAG2 reconstitution had virtually no effect on global TAD structure (Fig. S5), reinforcing prior observations that STAG1-cohesin is sufficient to maintain TAD architecture in the absence of STAG2 (6,22,23,24). Together, these data suggest that the primary consequence of STAG2 restoration is the regulation of chromatin loop size.

### Acute STAG2 reconstitution triggers a focused and primarily activating gene expression program

We next sought to determine if the reduction in chromatin loop size was accompanied by a corresponding shift in gene expression. To do this, we performed deep RNA-seq on biological triplicates of H4 and MOG-G-UVW cells 36 hours post-infection with adeno-STAG2 or vector control (Fig. 5A,B). Remarkably, the global reduction in loop scale did not trigger widespread gene expression changes. In H4 cells, using stringent criteria (p<0.05, FC>2), we identified only 78 differentially expressed genes (DEGs) out of 15,796 detected transcripts (<0.5%; Tables S2-S4). Consistent with these results, acute STAG2 reconstitution in MOG-G- UVW cells induced similarly modest global changes, with only 165 of 16,107 detected genes (∼1%) meeting the same threshold (p<0.05, FC>2; Tables S5-7). In both cell lines, the majority of DEGs (81% in H4; 68% in MOG-G-UVW) were upregulated, suggesting that STAG2-containing cohesin functions primarily as a transcriptional activator in this context.

**Figure 5.**
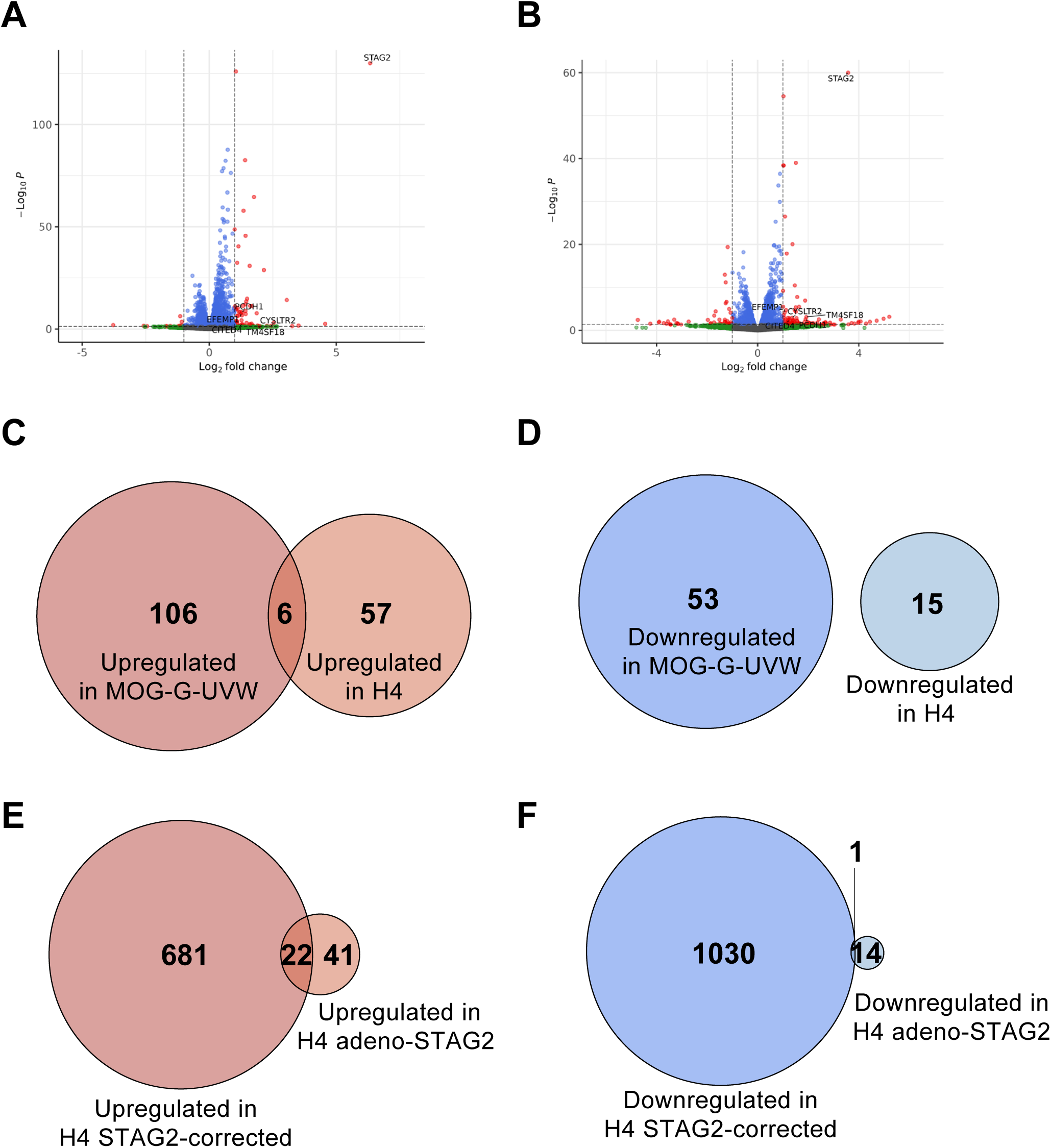
Impact of adenoviral STAG2 reconstitution on gene expression. Volcano plot of RNA-seq data comparing (A) H4 and (B) MOG-G-UVW cells infected with adeno-STAG2 versus vector alone (36 h post-infection). Data points are colored by significance: green, FC>2; blue, *p*<0.05; red, both FC>2 and *p*<0.05. *C*-*D*, Venn diagrams showing the overlap of significantly upregulated (C) and downregulated (D) genes between adeno-STAG2-infected H4 cells and adeno-STAG2-infected MOG-G-UVW cells. *E*-*F*, Venn diagrams showing the overlap of significantly upregulated (E) and downregulated (F) genes between adeno-STAG2-infected H4 cells and the permanently STAG2 corrected H4 88-1 cell line.

Gene Ontology (GO) analysis of the most highly altered genes (FC>2) in both lines yielded no significant pathway enrichment. However, applying a more inclusive threshold (FC>1.5) revealed broader regulatory signatures, including the canonical Wnt signaling pathway and positive regulation of gene expression (Fig. S6; Tables S8, S9). To identify the most robust targets of STAG2, we cross-referenced the DEGs from both models. Excluding STAG2 itself, five genes - CITED4, PCDH1, EFEMP1, CYSLTR2, and TM4SF18 - were upregulated in both MOG-G-UVW and H4 lines (Fig. 5C,D; Table S10).

Next, we compared the acute H4 adeno-STAG2 RNA-seq dataset to our previously published expression profile of permanently gene-corrected H4 cells (22). We observed that 35% (22/63) of the acutely upregulated genes in adeno-STAG2 infected H4 cells were also induced in permanently STAG2-corrected H4 cells (Fig. 5E). Notably, the magnitude of induction for these shared genes was generally higher in the STAG2-corrected H4 cells (Table S11), suggesting that STAG2-mediated activation at these loci intensifies over time. Conversely, only 3% (22/703) of the upregulated genes in the permanent model were also upregulated in our acute dataset. This limited overlap confirms that most expression changes in the permanently corrected cells represent secondary, indirect effects that accumulate long after STAG2 correction. Furthermore, the presence of only a single conserved downregulated gene (Fig. 5F, Table S11) reinforces the observation that direct transcriptional activation, rather than repression, is the primary consequence of STAG2 restoration.

### Kinetics of STAG2-mediated gene induction

To characterize the temporal dynamics of STAG2-dependent gene regulation, we used qRT-PCR to define the induction kinetics of target genes identified in our RNA-seq datasets. We prioritized 24 genes that were either shared between our acute and permanent H4 models (n=22) or conserved across both H4 and MOG-G-UVW lines (n=5). Three genes - CITED4, PCDH1, and EFEMP1 - were common to both groups. Following adeno-STAG2 transduction in H4 cells, STAG2 mRNA and protein levels peaked at 48 hours post-infection and declined rapidly thereafter (Fig. 6A,B).

**Figure 6.**
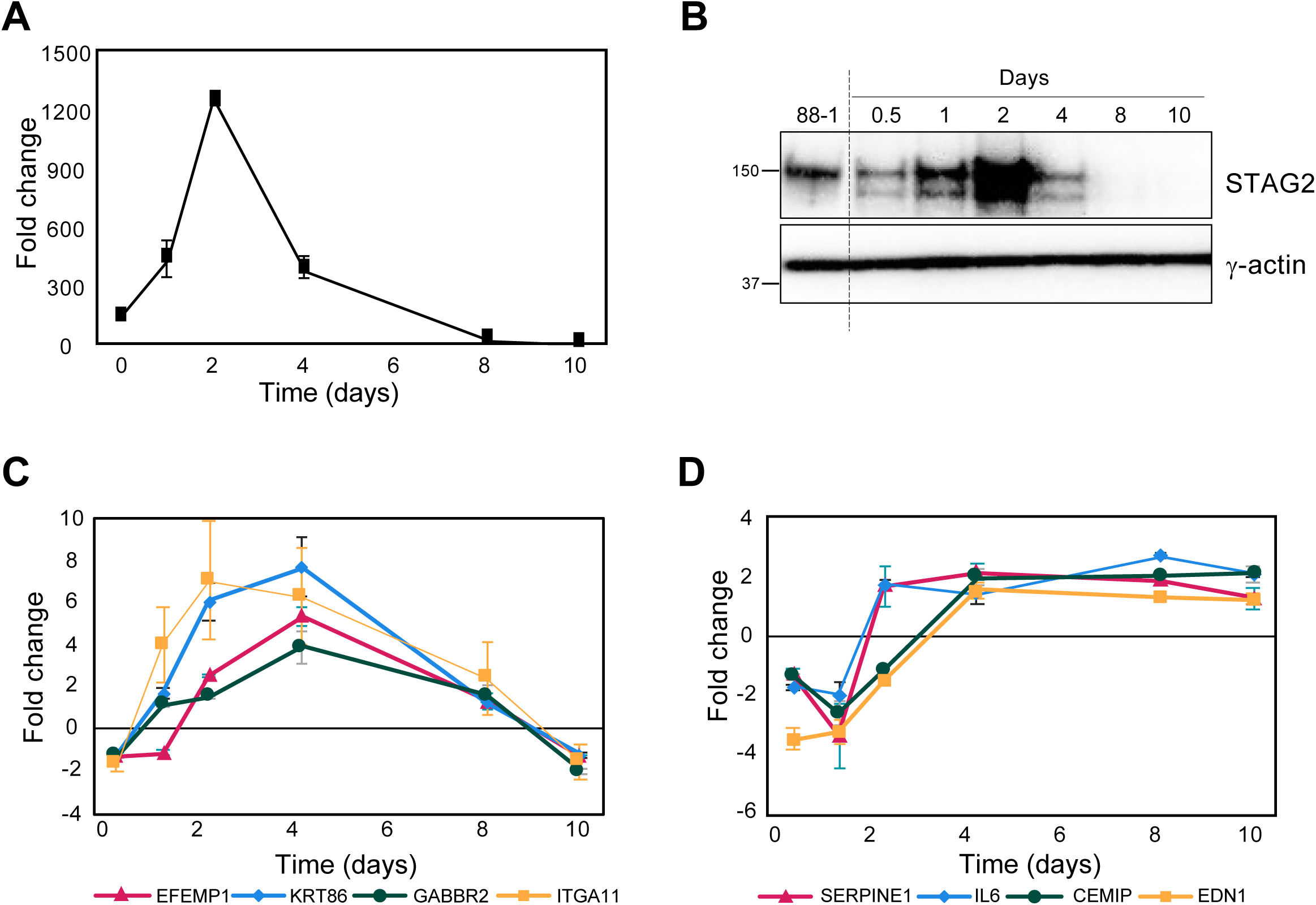
Induction kinetics of STAG2-regulated genes. *A*-*B*, Kinetics of STAG2 mRNA (A) and protein (B) expression in H4 cells at indicated time points following adeno-STAG2 infection, as determined by qRT-PCR and Western blot, respectively. *C*-*D*, qRT-PCR analysis of STAG2-induced genes over the same time course. Panel C illustrates representative genes requiring sustained STAG2 levels for continued expression, while panel D displays genes whose induction is maintained independently of STAG2 levels. Fold change refers to the increase in expression after infection with adeno-STAG2 compared to cells infected with adenoviral vector alone. Error bars represent S.D.

The induction profiles of the STAG2-responsive genes followed two distinct patterns. The expression of 16 genes closely mirrored the kinetics of STAG2, reaching maximal levels at 48 hours before diminishing (Fig. 6C, Fig. S7). In contrast, the remaining 7 genes remained elevated even after STAG2 expression returned to baseline (Fig. 6D, Fig. S7). We considered the first group as the most probable direct transcriptional targets of STAG2, as their sustained upregulation required the continued presence of STAG2 protein. Notably, EFEMP1 was the only gene in this high-confidence group that was upregulated by STAG2 in every experimental model tested.

### Conserved STAG2-responsive genes reside within stable, high-intensity chromatin loops

We next sought to define the structural environment of the five core STAG2-responsive genes that were upregulated in both H4 and MOG-G-UVW cells upon acute STAG2 reconstitution (CITED4, PCDH1, EFEMP1, CYSLTR2, and TM4SF18). Interestingly, high-resolution Hi-C revealed that each of these genes is situated within an exceptionally high-intensity chromatin loop (Fig. 7A-B, Figs. S8). Notably, at the level of resolution and sensitivity of our Hi-C analysis, the strength and anchors of these large-scale loops remained unchanged following STAG2 reconstitution (Figs. S9-10). This indicates that STAG2- responsive genes are unlikely to be regulated through the de novo formation of chromatin loops, but are instead positioned within pre-existing, structural stable loops.

**Figure 7.**
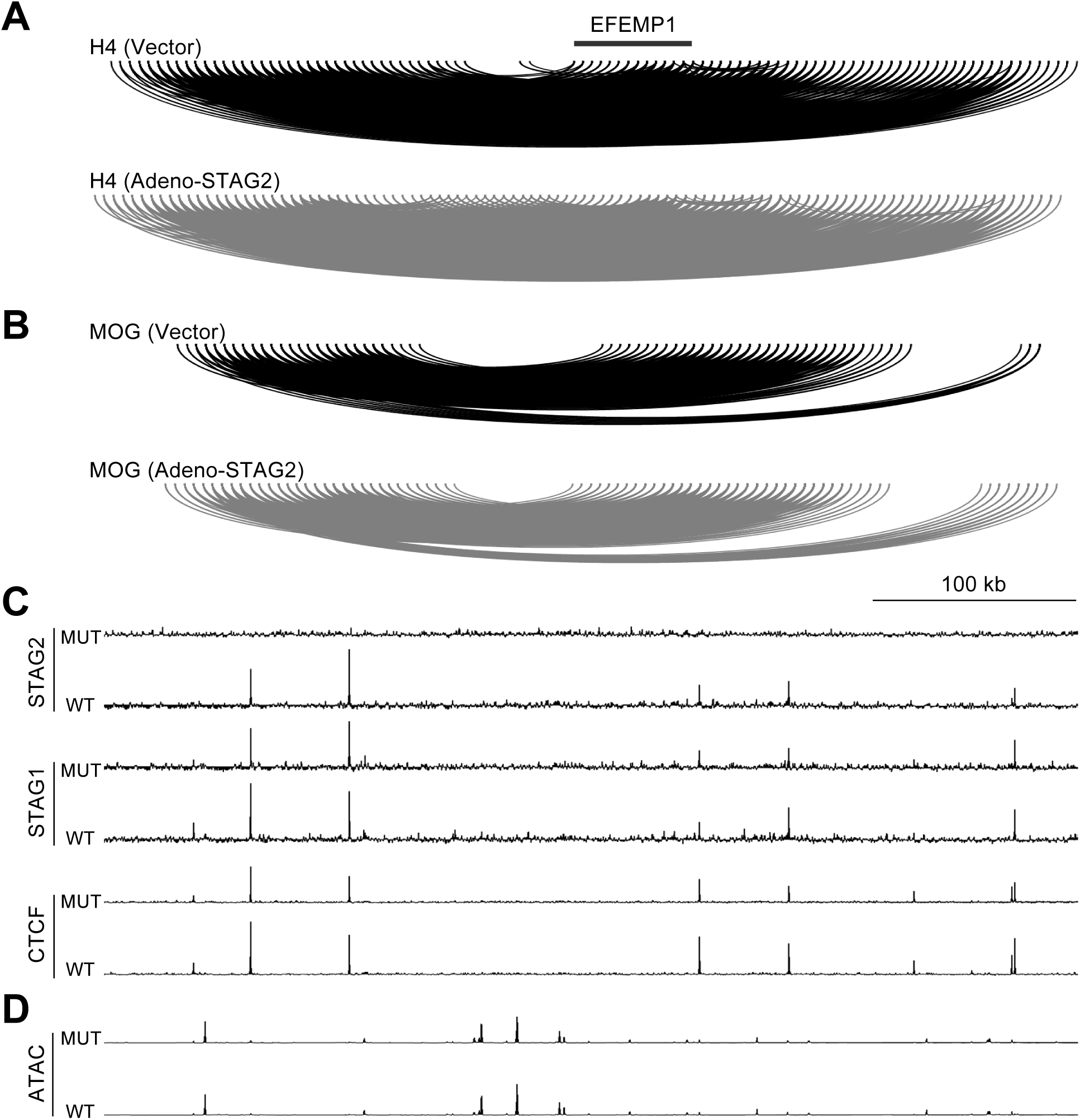
Integrative genomics surrounding the EFEMP1 locus. The depicted 500 kb region on chromosome 2 (hg38) shows the regulatory landscape of EFEMP1. *A-B*, Schematic of chromatin loop locations and intensities in STAG2-deficient H4 cells (A) and MOG-G-UVW cells (B) 36 hours after infection with adeno-STAG2 (WT) or vector control (MUT). Loop intensity heatmaps and quantification are provided in Fig. S9. *C*, ChIP-seq tracks for STAG2, STAG1, and CTCF in H4 cells (MUT) and STAG2-corrected 88-1 derivatives (WT). *D*, ATAC-seq profiles showing regions of open and closed chromatin in H4 cells infected with adeno-STAG2 (WT) or vector alone (MUT).

To further annotate this landscape, we performed ChIP-seq for STAG2, STAG1, and CTCF, alongside ATAC-seq to assess chromatin accessibility. While each of the five loci contained binding sites for all three cohesin-CTCF components, their distribution and local chromatin accessibility were unaffected by STAG2 status (Fig. 7C-D, Fig. S8). Collectively, these findings suggest that STAG2-mediated transcriptional activation is not driven by large-scale chromatin loop remodeling but instead likely results from the fine-tuning of dynamics within existing high intensity loops.

### EFEMP1 is a conserved growth-suppressive target of STAG2 in GBM

To identify high-priority STAG2 targets, we focused on genes consistently regulated across three distinct contexts: (i) acute reconstitution in H4 cells, (ii) acute reconstitution in MOG-G-UVW cells, and (iii) permanent genetic correction in H4 cells. Only three genes - EFEMP1, PCDH1, and CITED4 - met these stringent criteria. We further cross-referenced these candidates with transcriptomic data from a fourth isogenic system, permanently STAG2 corrected 42MGBA human GBM cells (1). Of these, only EFEMP1 was robustly regulated across all four models, showing a dramatic 28-fold induction upon STAG2 correction in 42MGBA cells. Given that EFEMP1 (Fibulin-3) is a secreted extracellular matrix protein with known inhibitory effects on GBM growth and invasion (25,26,27,28), we focused on it as a primary, conserved, putative STAG2 effector.

Because EFEMP1 is a secreted protein, we used ELISA to determine if these transcriptional changes translated to increased protein levels in the extracellular space. In both acute and permanent models, STAG2 restoration consistently drove the secretion of EFEMP1 protein at magnitudes comparable to the observed mRNA induction (Fig. 8A,B). We next investigated the regulatory elements governing this response using luciferase reporters containing regions of the EFEMP1 proximal promoter (1.9 kb and 2.3 kb). Notably, these reporters failed to recapitulate STAG2-responsiveness, indicating that the induction of EFEMP1 is not a localized promoter event, but instead likely depends on distal regulatory elements (Fig. S11).

**Figure 8.**
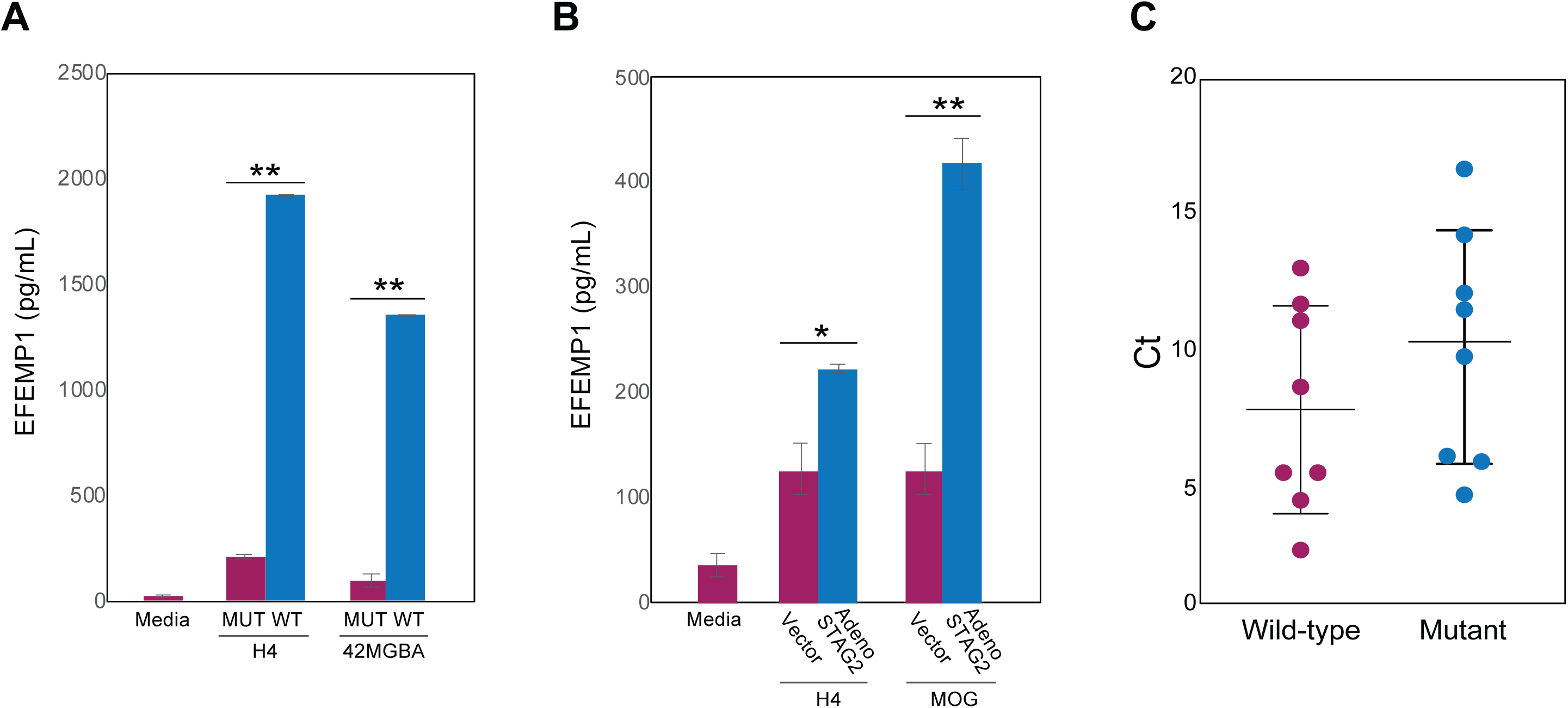
STAG2 regulates EFEMP1 expression and protein levels. *A,* Media conditioned for 24 hr. was collected from H4 cells, 42MGBA cells, and their permanently STAG2-corrected derivatives 88-1 and 53-1, respectively. EFEMP1 protein was measured by ELISA. *B,* Same as in (A) except for H4 and MOG-G-UVW cells infected with adeno-STAG2 or vector alone. All data are presented as mean ±SD. Statistical significance was determined by a Student’s t-test; *, p<0.05; **, p<0.01. *C,* qRT-PCR for *EFEMP1* and *GAPDH* (for normalization) in eight primary GBM xenografts harboring wild-type *STAG2* genes and eight harboring mutant *STAG2* genes. Lower normalized Ct values indicate higher levels of mRNA. Given the logarithmic nature of the Ct scale, the observed difference in means represents an approximately 6-fold difference in expression. Assays were performed in triplicate. Statistical significance was determined by an unpaired two-tailed t-test (p = 0.2770).

Finally, we explored the relationship between STAG2 mutational status and EFEMP1 expression in sixteen patient-derived GBM xenografts. We measured EFEMP1 mRNA levels via qRT-PCR in primary GBM xenografts with either wild-type (n=8) or mutant (n=8) STAG2 (29). Although the genetic heterogeneity across these genetically unrelated tumors precluded statistical significance in this cohort, STAG2 wild-type xenografts exhibited a 5.7-fold higher average EFEMP1 expression compared to their mutant counterparts (Fig. 8C). This trend provides supportive evidence that the regulation of EFEMP1 expression by STAG2 is a conserved feature of GBM biology.

## DISCUSSION

Current models of STAG2-mediated tumor suppression emphasize the role of the cohesin complex in shaping 3D genome architecture to regulate enhancer-promoter interactions. However, a major challenge in the field has been distinguishing the primary, direct consequences of STAG2 inactivation from the secondary, indirect changes that accumulate in steady-state models. By utilizing a recombinant adenovirus to rapidly restore physiological STAG2 levels to STAG2 mutant human cancer cells, we have identified early effects of STAG2 restoration - a genome-wide reduction in chromatin loop size, and the induction of a small number of target genes.

We find that rapid STAG2 reconstitution triggers an immediate reduction in chromatin loop size. This observation is consistent with prior biochemical findings that STAG2-cohesin possesses shorter chromatin residence times and lower loop extrusion processivity than STAG1-containing complexes (18). This observation is also consistent with recent findings in other malignancies, such as bladder cancer and acute myeloid leukemia, in which STAG2 loss has been associated with the formation of enlarged chromatin loops (23,30). We propose that STAG2-inactivating mutations promote oncogenesis by alleviating a restriction on the size of chromatin loops, allowing for an expanded range of chromatin interactions that disrupts the maintenance of a tumor-suppressive transcriptome.

Despite this global alteration in chromatin loop size, fewer than 1% of genes exhibited significant changes in expression. STAG2-induced genes appear to reside within stable, high-intensity loops. This suggests that STAG2-mediated transcriptional regulation may occur through subtle changes in relatively stable chromatin loops that are not captured by steady-state Hi-C.

Among these targets, we identify EFEMP1 (fibulin-3) as the most robust putative STAG2 effector (25). EFEMP1 was consistently induced across all four GBM models tested, with induction levels reaching 28-fold in permanent correction models. We confirmed that this transcriptional activation leads to increased secretion of EFEMP1 protein into the extracellular matrix. Given that EFEMP1 is known to influence tissue structure and cell signaling and has previously been reported to have tumor-suppressive roles in GBM (26,27,28), its regulation by STAG2 could be central to the STAG2 mutant phenotype. However, confirming whether EFEMP1 is a mediator of STAG2 function will require further testing in in vivo models.

While our data demonstrates that acute modulation of STAG2 leads to specific alterations in chromatin loop size and gene expression, we acknowledge that there are additional mechanisms that may contribute to its tumor suppressive activity. For example, STAG2 inactivation may alleviate a putative ‘caretaker’ function (31). It is also possible that STAG2 mutations function indirectly to modulate the activity of co-occurring oncogenic mutations. This model is consistent with findings in Ewing’s sarcoma, where STAG2 loss has been shown to modulate the oncogenic program of the EWS-FLI fusion protein (32).

Furthermore, our acute restoration experiments may not fully reverse the epigenetic reprogramming established during the long period of STAG2 deficiency. It is also possible that reconstituting chromatin-remodeling proteins may require more time to manifest than reconstituting proteins involved in signaling pathways. STAG2 inactivation triggers a shift toward an exclusively STAG1-cohesin state, which likely possesses oncogenic properties. However, the transition away from this STAG1-dominant state could be gradual.

In summary, this study provides a high-resolution view of the primary effects of STAG2 reconstitution in mutant human cancer cells. Our findings indicate that STAG2 serves as a fundamental regulator of chromatin loop scale and identify EFEMP1 as a promising putative effector of STAG2 tumor suppression in GBM.

## EXPERIMENTAL PROCEDURES

### Generation of a STAG2-Expressing Adenovirus

The STAG2-expressing adenovirus was assembled and packaged using the Ad-Easy system (details in ref. 15). Briefly, the wild-type STAG2 open reading frame (CCDS43990) was cloned into the pAdTrack-CMV shuttle vector. The insert was digested with BamHI/NotI and ligated into the BglII/NotI-digested vector. This STAG2-containing CMV-GFP shuttle vector was then linearized with PmeI and electroporated into electrocompetent BJ5183-AD-1 *E. coli* (Agilent) containing the adenoviral backbone plasmid pAdEasy-1. Recombinants were identified by restriction digestion. Thr resulting adeno-STAG2 and empty vector control were then linearized with PacI and transfected into Ad-293 cells (Agilent) using Lipofectamine 3000 (Invitrogen). Virus was harvested ten days post-transfection via four consecutive freeze thaw cycles in a dry ice/ ethanol bath and a 37-degree water bath. These primary viral lysates were amplified through two serial passages in Ad-293 cells to achieve final viral titers of approximately 1×10^6^ infectious particles/ul, as determined by GFP flow cytometry.

### Cell Lines

The H4 human GBM cell line was purchased from the American Type Culture Collection (ATCC). The MOG-G-UVW cell line was purchased from the European Collection of Authenticated Cell Cultures (ECACC). 42MGBA cells were purchased from the German Collection of Microorganisms and Cell Cultures (DSMZ). The isogenic, gene edited derivatives of H4 and 42MGBA cells (designated 88-1 and 53-1, respectively) were generated as previously described (1). All cell lines were maintained in Dulbecco’s modified Eagle’s medium (DMEM) supplemented with 10% fetal bovine serum and 1% penicillin/streptomycin at 37 °C in 5% CO2. Cell lines were authenticated by short tandem repeat (STR) profiling and routinely tested for mycoplasma contamination.

### Western Blotting

Total protein lysates were prepared in RIPA buffer (50 mM Tris-HCl, pH 8.0, 150 mM NaCl, 1.0% NP-40, 0.5% sodium deoxycholate, and 0.1% SDS). Protein concentrations were determined using a BCA assay (Pierce). Equivalent amounts of protein (20-40 µg per lane) were dissolved in 1X NuPAGE LDS sample buffer, boiled for 5 min, and separated by SDS-PAGE. Proteins were electro-transferred onto polyvinylidene difluoride (PVDF) membranes (Millipore), which were then blocked in 5% (w/v) non-fat dry milk in TBST (Tris-buffered saline with 0.1% Tween-20) for 1 h at room temperature. Membranes were incubated with a 1:1000 dilution of primary antibody against STAG2 (Santa Cruz Biotechnology, Cat. No. sc-81852) with gentle agitation overnight at 4°C. After incubation with the appropriate horseradish peroxidase-conjugated secondary antibody (Cell Signaling Technology) for 1 h at room temperature, membranes were developed using SuperSignal West Pico PLUS chemiluminescent substrate (Pierce) and imaged on a myECL Imager.

### Hi-C Library Preparation, Sequencing, and Bioinformatic Analysis

Hi-C libraries were generated using the Arima Hi-C+ kit (Arima Genomics) according to the manufacturer’s instructions. Libraries were sequenced on an Illumina NovaSeq 6000 instrument to generate paired-end sequencing data. Full-length paired-end reads were aligned to the hg19 reference human genome using parallelized BWA (33). Next, the resulting two SAM files for each end were merged into a single file, and only uniquely mapped paired-end reads (∼60% of the total) were retained. Subsequently, redundant read pairs that shared the same chromosome and start sites were removed. After the removal of PCR duplicates, we assigned read pairs to GATC/GANTC-digested fragments and discarded read pairs located within the same fragment. Next, based on strand information, we assigned the remaining reads to three categories - inward, outward and same-strand. We kept “inward” read pairs if the distance between two fragments was >1kb, and “outward” read pairs if the distance was >5kb. Filtered “inward”, “outward” and all “same-strand” were then merged together as cis read pairs. After these filtering steps, we obtained cis and trans fragment pairs. Since the analysis pipeline is conducted at 5 kb resolution, we assigned each fragment pair to 5kb anchor pairs. We then applied the *HiCorr* bias correction algorithm to the data as described (19) followed by the LoopEnhance tool from *DeepLoop* to enhance low depth data and remove noisy pixels (20). The enhanced loop strength was normalized by dividing by the value of the 300,000th-ranked loops. For all pairwise comparisons between STAG2 wild-type and STAG2-mutant samples, we merged the top 300k loops from each condition, generating a total of approximately 380k loops. We then compared the strength of these loops in STAG2-mutant and STAG2-corrected samples. Using a cutoff of fold change > 2, we identified lost, gained and common loops.

### RNA-seq Library Preparation, Sequencing, and Transcriptomic Analysis

Total RNA was isolated using the RNeasy Mini Kit (Qiagen) according to the manufacturer’s instructions. RNA-seq libraries were prepared using the NEBNext Ultra II RNA Library Prep Kit (New England Biolabs) and sequenced on an Illumina NovaSeq 6000 instrument to a target depth of approximately 150 million reads per sample. Raw sequencing reads were quality-filtered and aligned to the human reference genome (hg19) using HISAT2 (34). Gene-level read counts were quantified using featureCounts (35). The statistical tests for RNA-seq data, normalizing raw data and identifying DEGs in this study were derived from a statistical model from DESeq2 (36).

### Quantitative Real-Time PCR (qRT-PCR)

Total RNA was prepared by standard TRIZOL-based methods. Quantitative reverse transcription-PCR (qRT-PCR) was performed in StepOnePlus Real-Time PCR Systems using TaqMan gene expression assays (Applied Biosystems, Foster City, CA) and the Superscript III Platinum One Step qRT-PCR System (Invitrogen, Carlsbad, CA), according to the manufacturers’ specifications. Relative gene expression levels were calculated using the 2−Δ(ΔCT) method, normalizing to the expression of the GAPDH housekeeping gene. All assays were performed at least in triplicate.

### Chromatin Immunoprecipitation and Sequencing (ChIP-seq)

Cross-linked chromatin was prepared using the SimpleChIP® Plus Sonication Chromatin IP Kit (Cell Signaling Technology) according to the manufacturer’s protocol. ChIP-seq libraries were then prepared using the DNA Library Prep Kit for Illumina (Cell Signaling Technology). The resulting libraries were sequenced on an Illumina NovaSeq 6000 instrument.

### Assay for Transposase-Accessible Chromatin (ATAC-seq)

To assess chromatin accessibility, H4 cells infected with either adeno-STAG2 or empty vector were harvested for nuclei preparation. Isolated nuclei were subjected to a tagmentation reaction using Tn5 transposase (from the Active Motif ATAC-seq kit). The tagmented DNA was purified and amplified via limited-cycle PCR to generate libraries containing unique barcoded index sequences for multiplexing. The resulting libraries were sequenced on an Illumina NovaSeq 6000 instrument.

### EFEMP1 Enzyme-Linked Immunosorbent Assay (ELISA)

To quantify secreted protein levels, equivalent numbers of cells were plated and cultured to confluence. The conditioned media were then collected, centrifuged to clear cellular debris, and assessed for the presence of EFEMP1 using a human EFEMP1-specific ELISA (Abcam) according to the manufacturer’s instructions. Standard curves were generated using dilutions of purified EFEMP1 protein to quantify EFEMP1 levels within the samples. All measurements were performed in triplicate.

### Luciferase Reporter Assays

To investigate the regulatory elements governing EFEMP1 expression, luciferase reporter assays were conducted using the pGL4.23 minimal promoter-driven firefly luciferase vector (Promega). Specific EFEMP1 promoter regions were amplified by PCR and cloned into the pGL4.23 vector. MOG-G-UVW cells were seeded in 12-well plates and transfected with 500 ng of the pGL4.23-EFEMP1-promoter constructs and 50 ng pRL-TK (Renilla luciferase) as an internal control for transfection efficiency using X-tremeGENE-9 (Sigma-Aldrich). Twenty-four hours after transfection, cells were infected with either adeno-STAG2 or empty vector and incubated for an additional 24 hours. Luciferase activities were measured using the Dual-Luciferase Reporter Assay System (Promega) according to the manufacturer’s protocol. Firefly luciferase activity was normalized to Renilla luciferase activity for each sample.

## Data Access

All raw RNA-seq and H-C sequencing data were submitted to the Gene Expression Omnibus (GEO). Accession numbers GSE318550 and GSE318552. HiCorr and DeepLoop pipelines for chromatin loop identification are available at https://github.com/JinLabBioinfo.

## Data Availability

All data are contained within the article and its supporting information. Any additional raw data that support the findings of this study are available from the corresponding author upon request.

## Acknowledgements

We thank Ash Dorsey for mathematical modeling of adenovirus titers. Supported by grant R01CA267872 (TW and FJ), The Children’s Cancer Foundation (TW), and Hyundai Hope on Wheels (TW).

## AI Disclosure

During the preparation of this manuscript, the authors used Google Gemini for the purpose of editing the text to improve clarity, grammar, and structural flow. The authors subsequently reviewed and edited the content as necessary and assume full responsibility for the accuracy and integrity of the published work.

---

**Figure S1.**
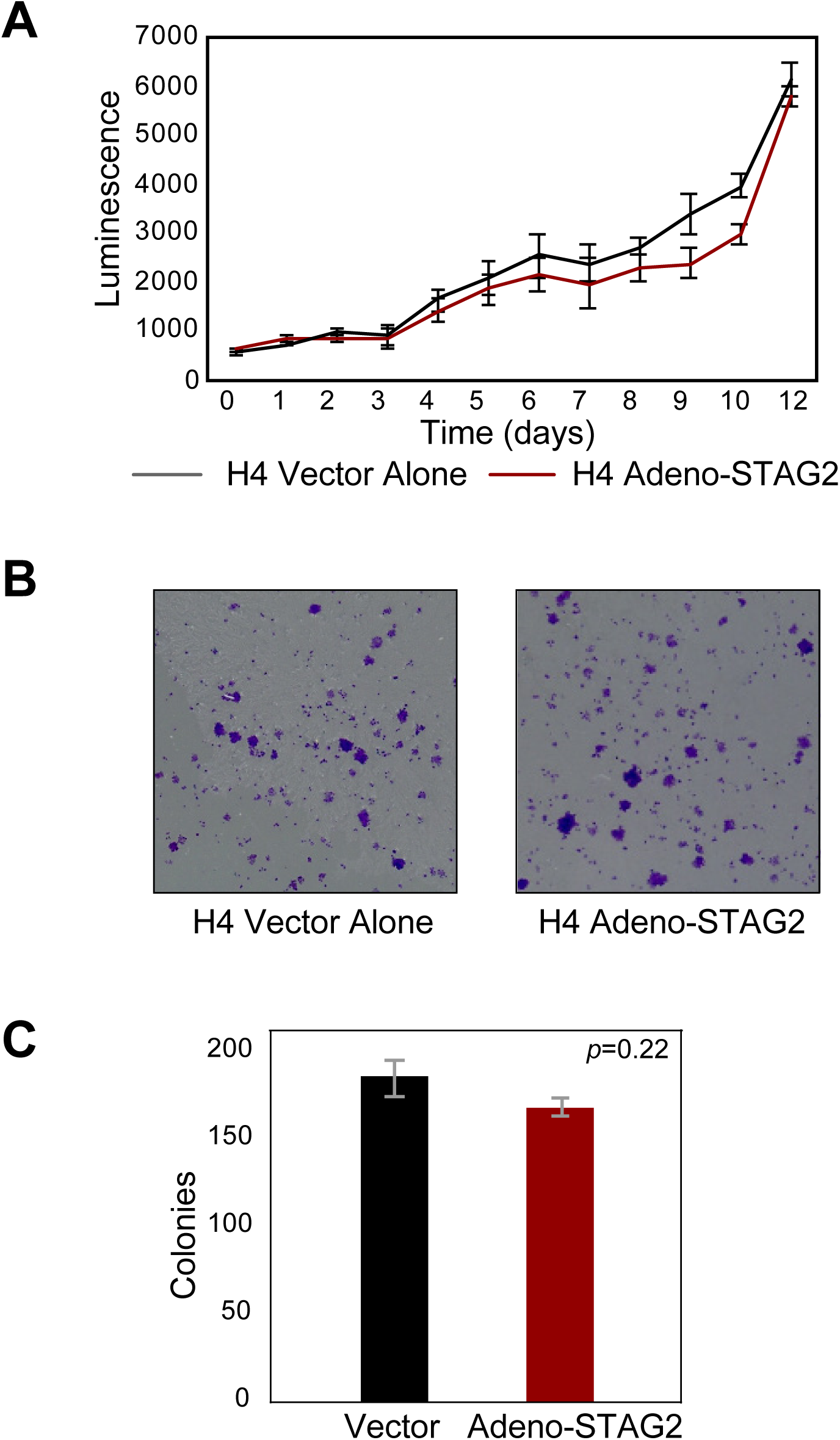
Impact of adenoviral STAG2 reconstitution on proliferation and colony formation. *A*, Proliferation of H4 cells infected with adeno-STAG2 or vector alone, measured using the CellTiter-Glo assay over 12 days. *B*,*C* Representative images and quantification of colony formation assays using the same cells as in (A). Error bars indicate S.D.

**Figure S2.**
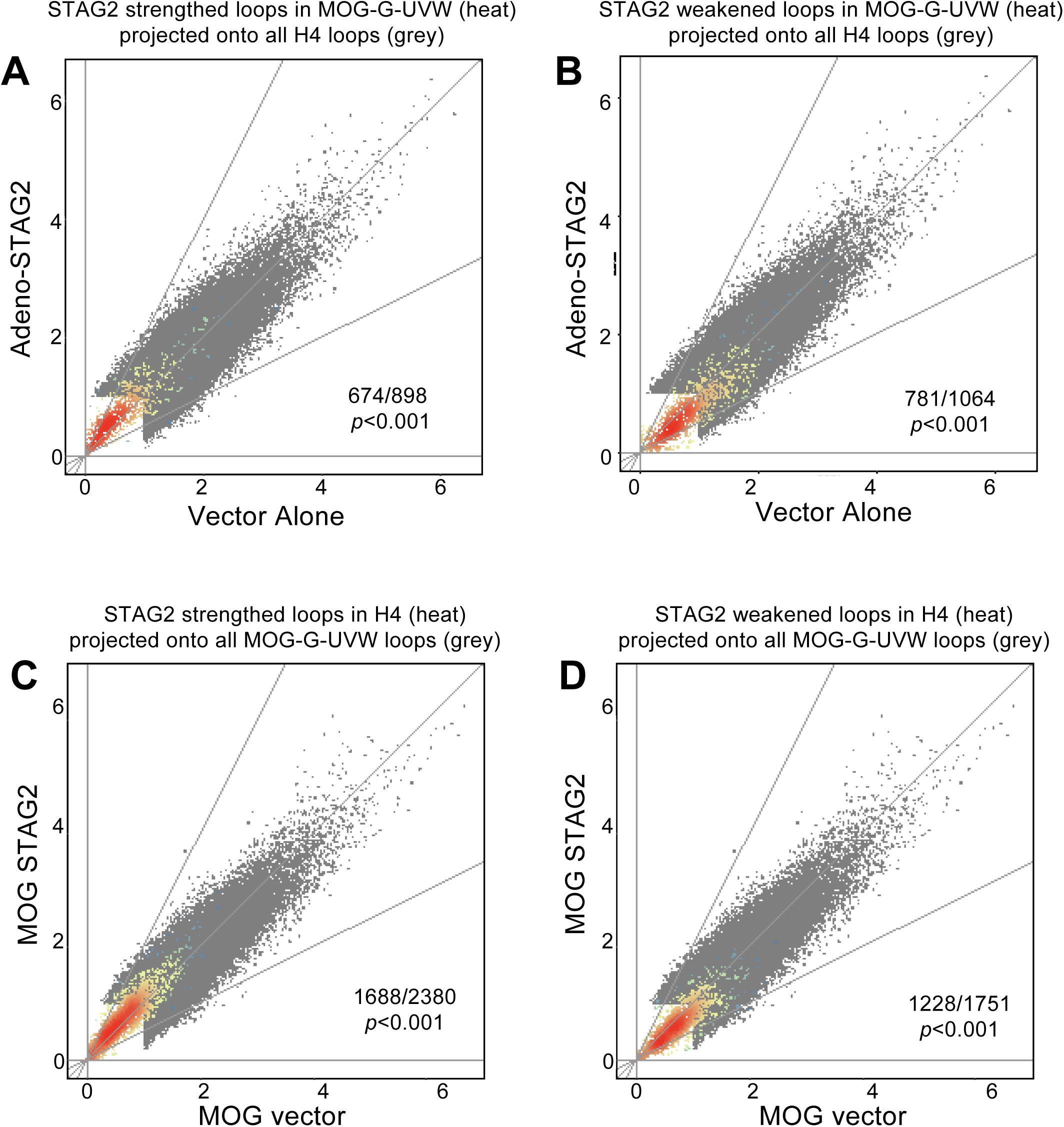
Conservation of individual STAG2-regulated loops across cell lines. A-D, Grey background scatter represents the intensity of the strongest ∼300,000 chromatin loops in H4 cells (A,B) and MOG-G-UVW cells (C,D) infected with adenoviral vector alone (x-axis) and adeno-STAG2 (y-axis). Heat (red/yellow) represents the projection of STAG2 strengthed (A and C) and STAG2 weakened loops (B and D) from the converse cell line onto the grey scatter. The fraction of the projected loops showing consistent regulation in the two cell lines is indicated together with a *p*-value.

**Figure S3.**
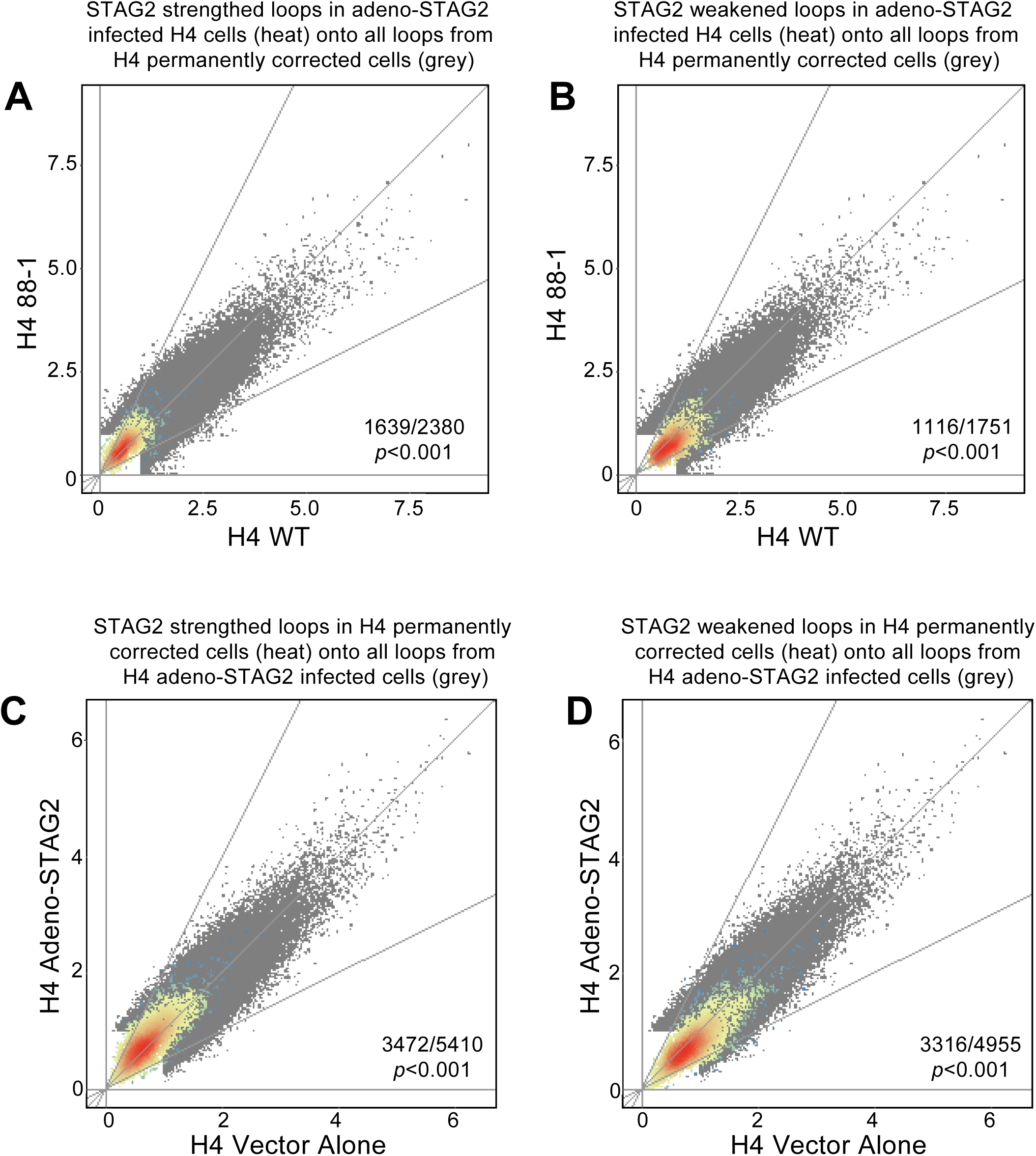
Conservation of individual STAG2-regulated loops in transiently and permanently STAG2-corrected H4 cells. A-D, Grey background scatter represents the intensity of the strongest ∼300,000 chromatin loops in permanently STAG2-corrected H4 cells (A,B) and adeno-STAG2 infected H4 cells (C,D). Heat (red/yellow) represents the projection of STAG2 strengthed (A and C) and STAG2 weakened loops (B and D) from the converse cell system onto the grey scatter. The fraction of the projected loops showing consistent regulation in the experimental systems is indicated together with a *p*-value.

**Figure S4.**
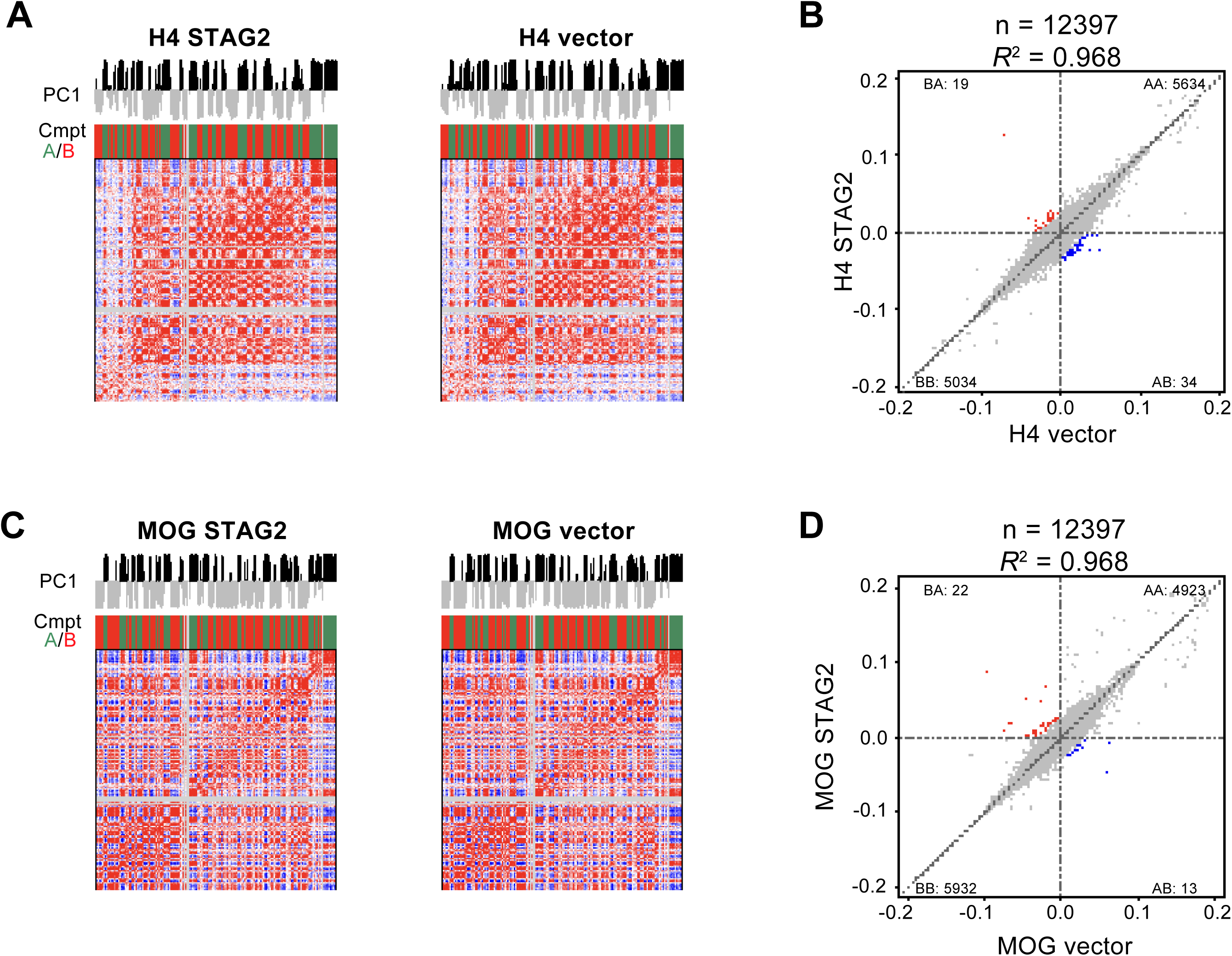
Impact of rapid STAG2 reconstitution on A/B compartment assignments in H4 and MOG-G-UVW cells. *A* and *C*, representative heatmaps of chromosome 2 showing Hi-C correlation matrices at 250-kb resolution for H4 cells (A) and MOG-G-UVW cells (C) infected with adeno-STAG2 or vector alone. Principal Component 1 (PC1) values define the A (green) or B (red) compartment status of individual 250-kb bins. *B* and *D*, scatterplot analysis depicting the effect of rapid STAG2 reconstitution on compartment assignments in H4 cells (B) and MOG-G-UVW cells (D). Bins undergoing compartment switches (p<0.01) are indicated in red (B to A) and blue (A to B). Coefficient of determination (*R*²) and the total number of bins (n) are indicated.

**Figure S5.**
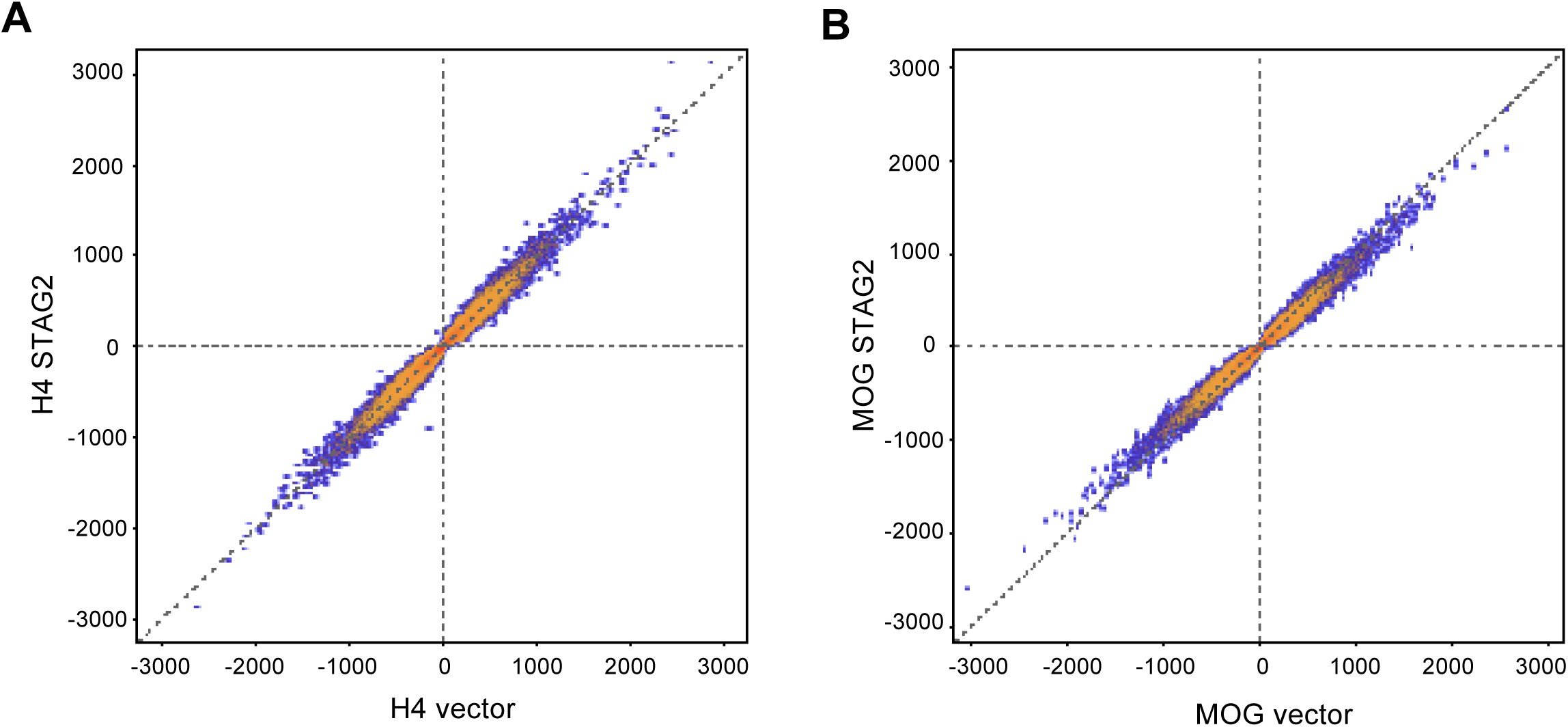
Impact of rapid STAG2 reconstitution on topologically associating domains (TADs) in H4 and MOG-G-UVW cells. Scatterplots depicting the effect of rapid STAG2 reconstitution on directionality indices (DI; a measure of TAD boundary strength) in H4 (A) and MOG-G-UVW (B) cells infected with adeno-STAG2 or vector alone.

**Figure S6.**
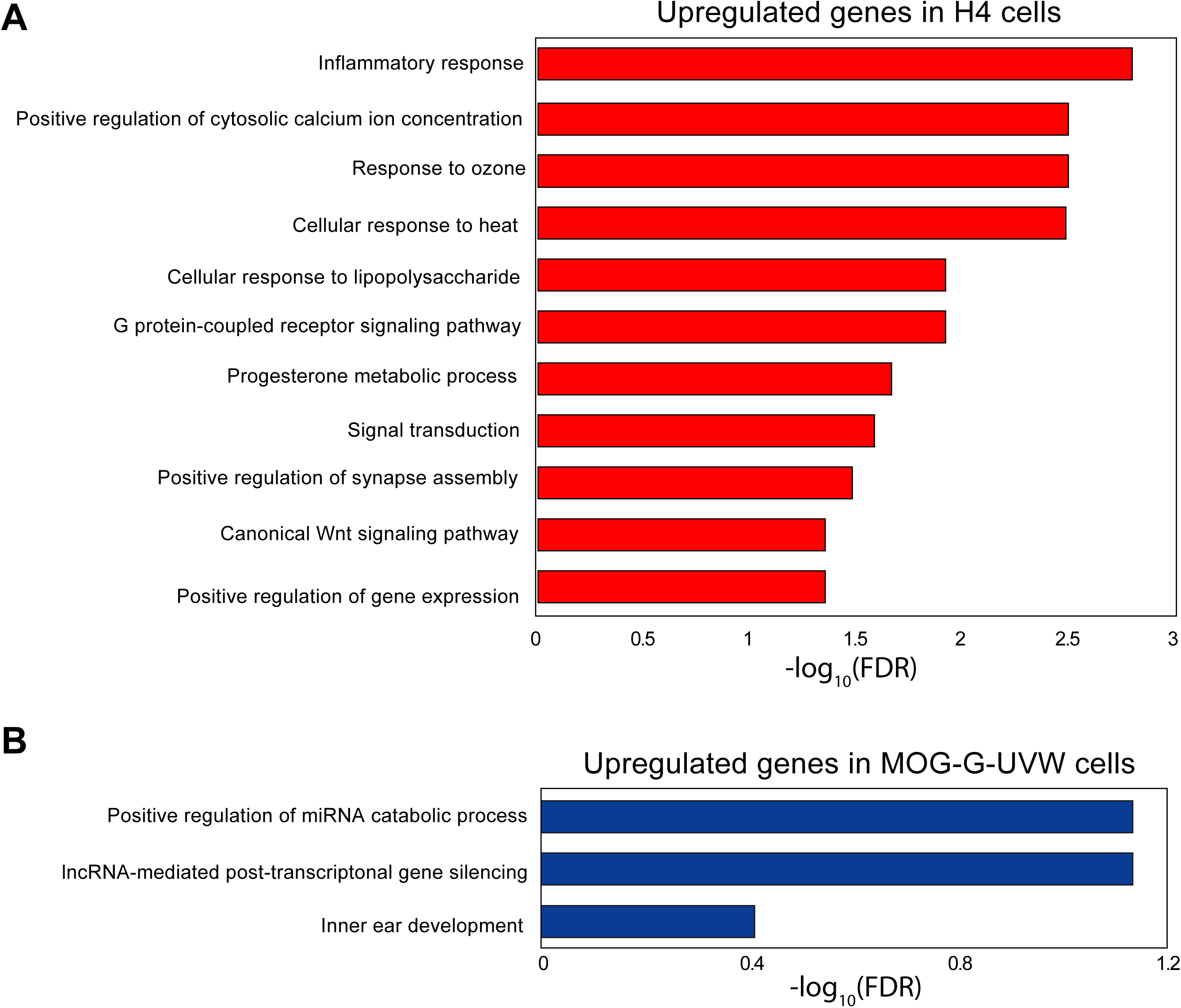
Gene Ontology (GO) analysis of STAG2-regulated genes. *A*, GO terms overrepresented in genes upregulated (FC>1.5) in H4 cells after infection with adeno-STAG2 (FDR<0.05). No GO terms overrepresented in downregulated genes met FDR<0.05 and therefore are not shown. *B*, Same as A for MOG-G-UVW cells.

**Figure S7.**
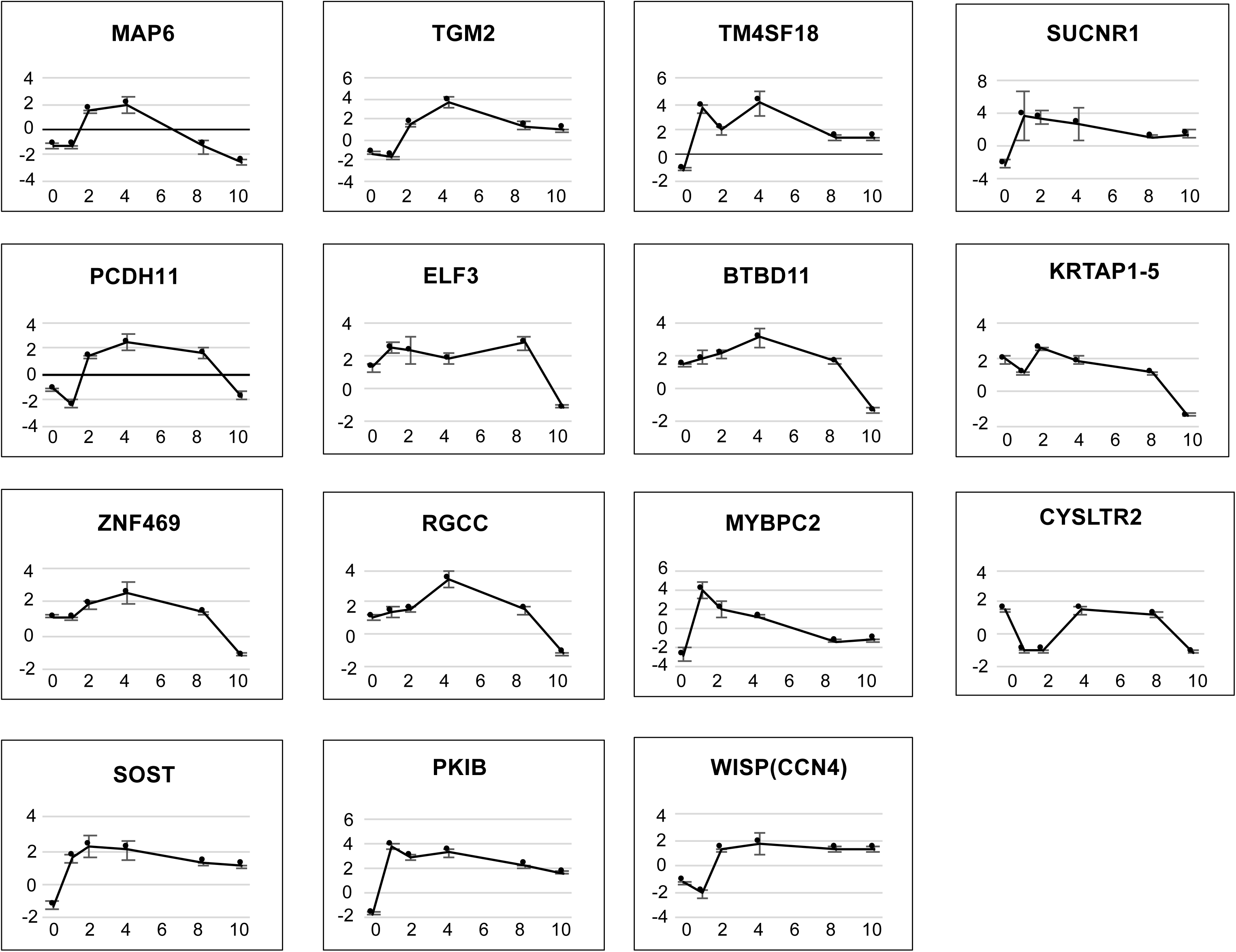
Induction kinetics of STAG2-regulated genes. Expression of STAG2 regulated genes at the indicated days following infection of H4 cells with adeno-STAG2 or vector alone, as measured by qRT-PCR. Fold change indicates the increase in expression after infection with adeno-STAG2 normalized to expression in cells infected with vector alone. Error bars represent S.D.

**Figure S8.**
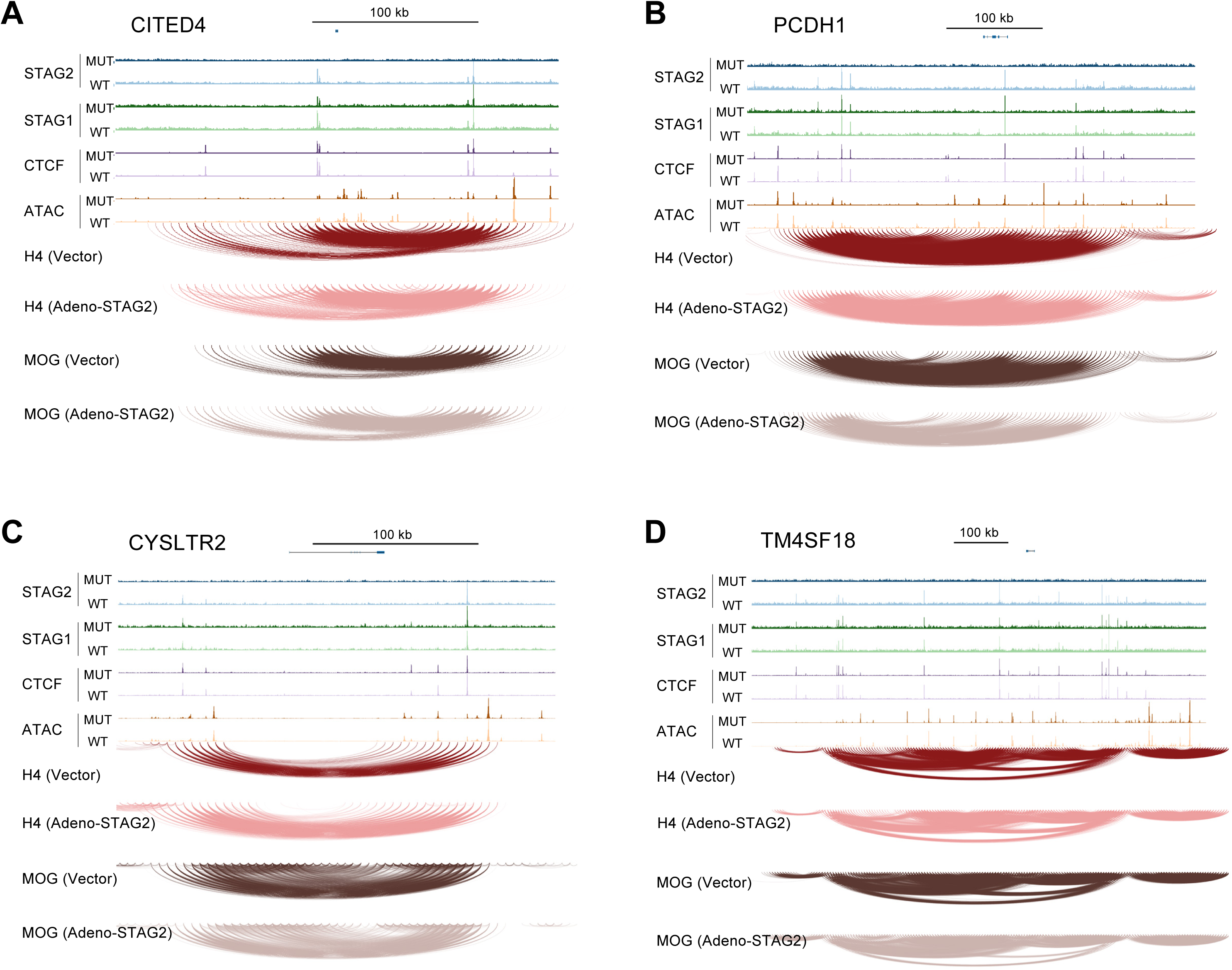
Integrative genomics surrounding the CITED4, PCDH1, CYSLTR2, and TM4SF18 loci. Schematic representation of the location and intensity of STAG2, STAG1, and CTCF chromatin binding sites by ChIP-seq (top); regions of open and closed chromatin by ATAC-seq (middle); and chromatin loops (bottom) surrounding the (*A*) CITED4 (chr1:41,180,006–41,474,755), (*B*) PCDH1 (chr5:140,963,663–141,476,413), (*C*) CYSLTR2 (chr13:49,111,678–49,401,750), and (*D*) TM4SF18 (chr3:148,489,014–149,408,708) loci.

**Figure S9.**
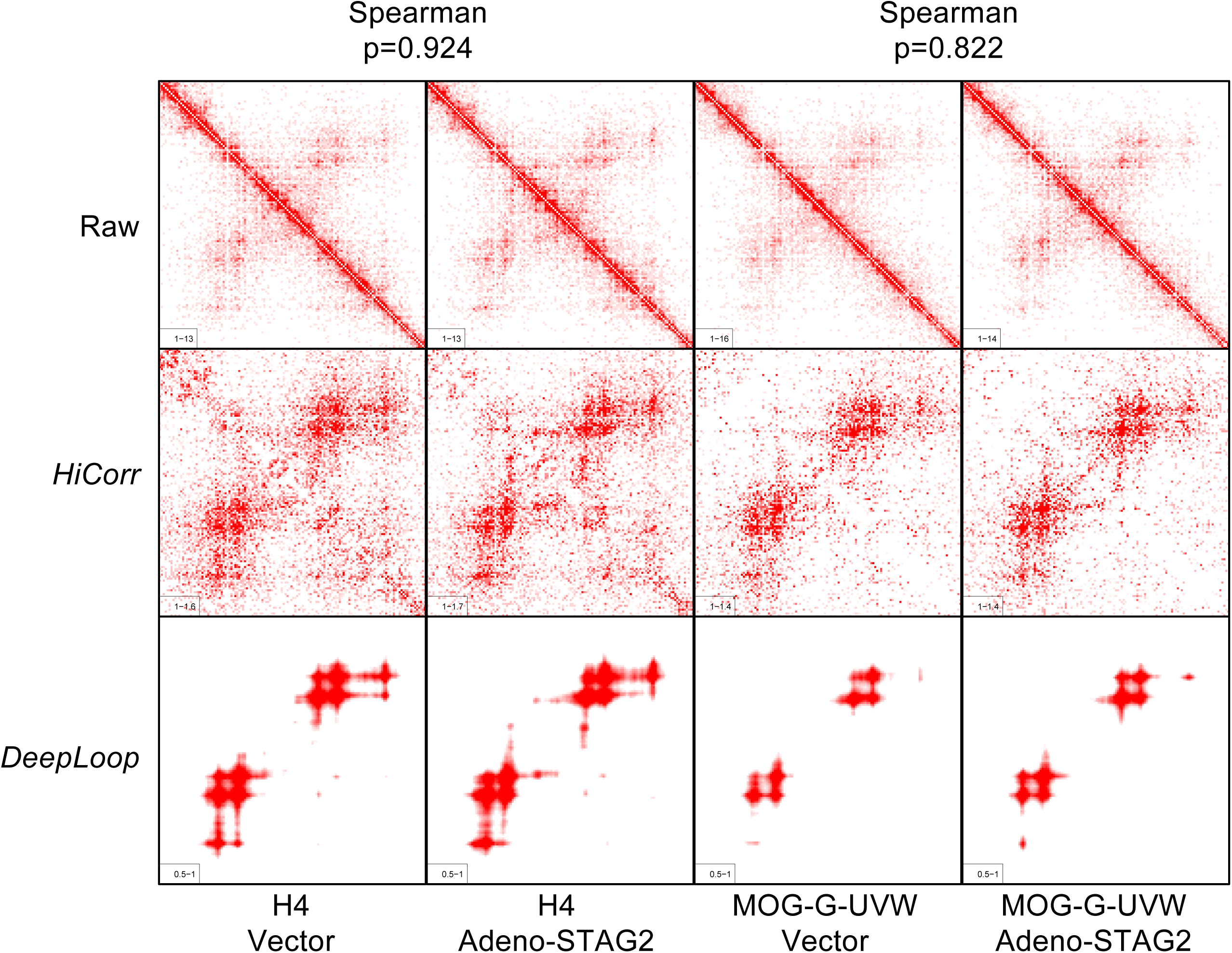
EFEMP1 locus Hi-C contact maps. Representative heatmaps showing normalized loop intensity at the EFEMP1 locus (chr2:55,797,886–56,456,442) in H4 and MOG-G-UVW cells. Data are presented for cells 36 hours post-infection with either adeno-STAG2 or an empty vector control. Maps are shown at 5-kb resolution following noise correction (HiCorr) and enhancement (DeepLoop). High Spearman correlation coefficients between adeno-STAG2 and vector-only conditions (0.924 for H4; 0.822 for MOG-G-UVW) indicate that chromatin loop structure and intensity surrounding EFEMP1 remain largely unchanged by STAG2 expression.

**Figure S10.**
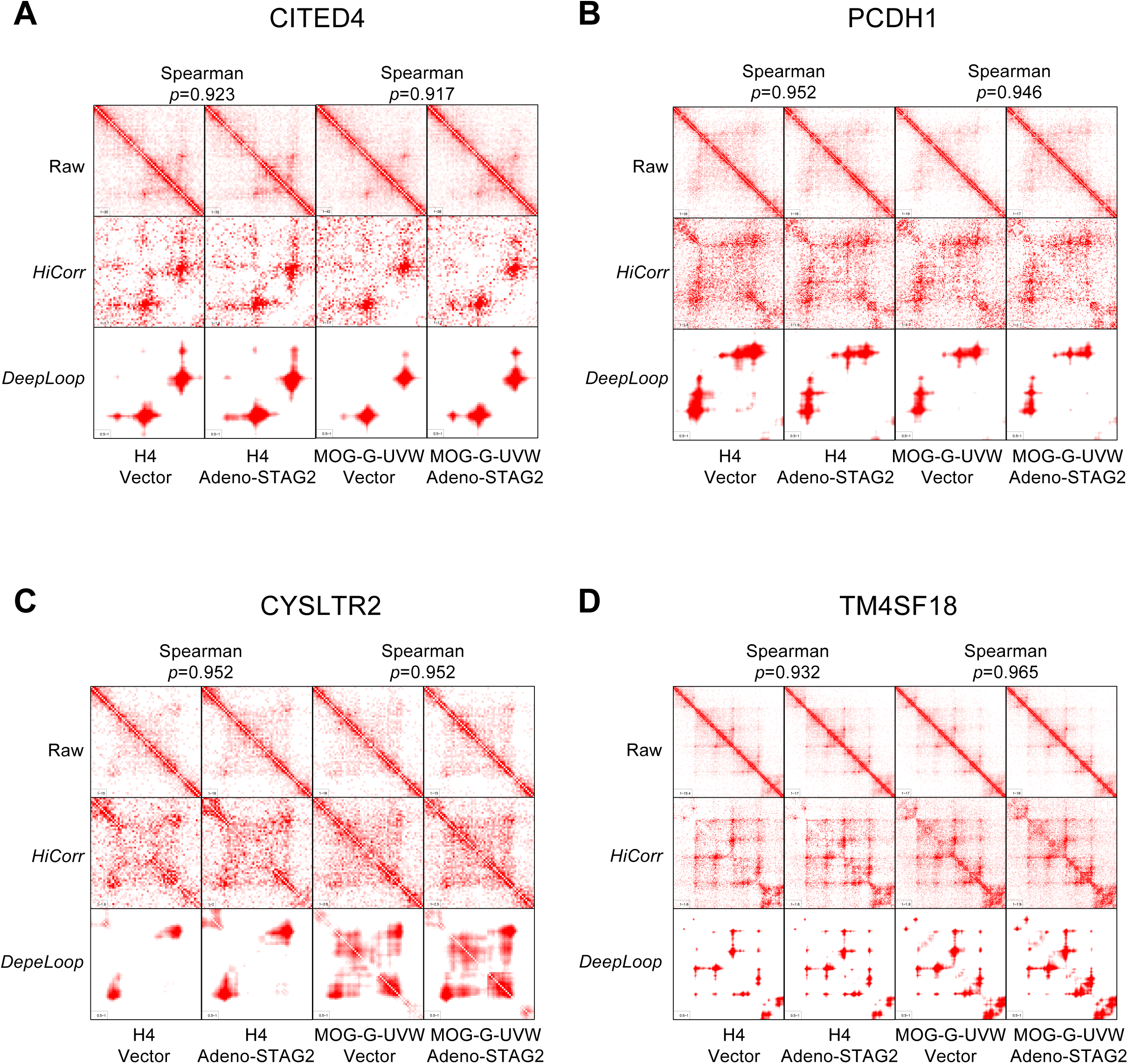
Hi-C contact maps for the CITED4, PCDH1, CYSLTR2, and TM4SF18 loci. Representative heatmaps showing normalized loop intensity at 5-kb resolution for four genomic regions: CITED4 (chr1:41,180,006–41,474,755), PCDH1 (chr5:140,963,663–141,476,413), CYSLTR2 (chr13:49,111,678–49,401,750), and TM4SF18 (chr3:148,489,014–149,408,708). Data are shown for H4 and MOG-G-UVW cells 36 hours post-infection with adeno-STAG2 or an empty vector control. Maps were processed via noise correction (HiCorr) and enhancement (DeepLoop). High Spearman correlation coefficients across all regions in both H4 and MOG-G-UVW lines (e.g., 0.924 for H4; 0.822 for MOG-G-UVW) indicate that chromatin architecture and loop intensity remain largely unchanged by STAG2 expression.

**Figure S11.**
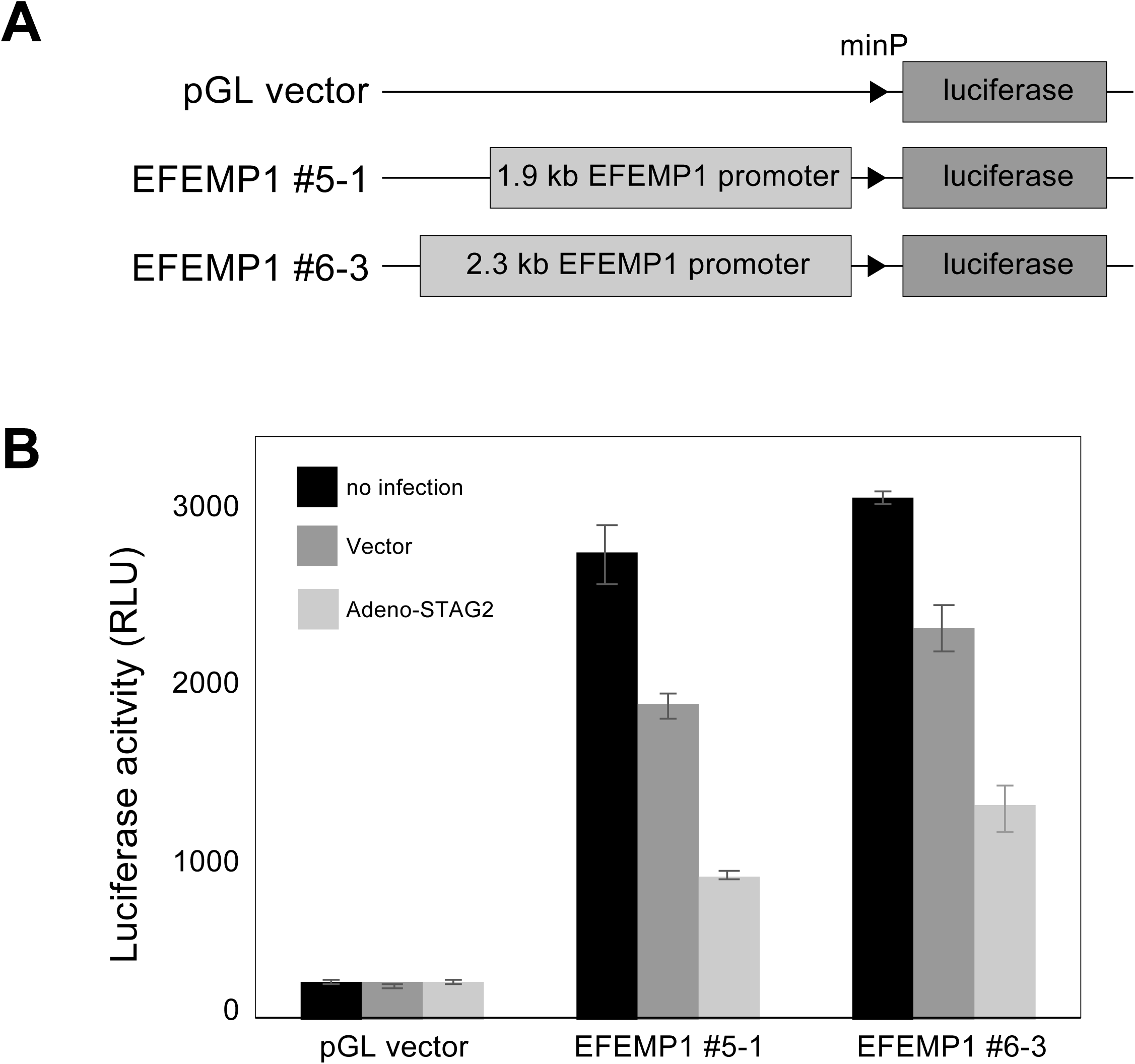
EFEMP1 luciferase reporter assay. *A*, Schematic representation of luciferase reporter constructs driven by the EFEMP1 promoter. *B*, MOG-G-UVW cells were transfected with the EFEMP1 promoter-luciferase reporter constructs and subsequently infected with either adeno-STAG2 or an empty vector control. Luciferase activity was measured 24 hours post-infection and normalized to Renilla luciferase activity. Data represent the mean ± S.D.

